# The host exocyst complex is targeted by a conserved bacterial type III effector protein that promotes virulence

**DOI:** 10.1101/2020.11.06.371260

**Authors:** Vassiliki A. Michalopoulou, Konstantinos Kotsaridis, Glykeria Mermigka, Dina Kotsifaki, Michael Kokkinidis, Patrick H. N. Celie, Jonathan D.G. Jones, Panagiotis F. Sarris

**Affiliations:** Department of Biology, University of Crete, 714 09 Heraklion, Crete, Greece; Institute of Molecular Biology and Biotechnology, Foundation for Research and Technology-Hellas, Heraklion, 70013, Crete, Greece; Division of Biochemistry, the Netherlands Cancer Institute, Amsterdam, the Netherlands; The Sainsbury Laboratory, Norwich Research Park, Norwich, United Kingdom; Biosciences, University of Exeter, Exeter, United Kingdom

## Abstract

For most Gram-negative bacteria, pathogenicity largely depends on the type-III secretion system that delivers virulence effectors into eukaryotic cells. The subcellular targets for the majority of these effectors remain unknown. Here, we show that *Xanthomonas campestris*, the causal agent of black rot disease, delivers the highly conserved effector XopP that interacts with host’s EXO70 protein. EXO70 is an essential component of the exocyst complex with a role in plant immunity. The XopP/EXO70 interaction is specific and inhibits exocyst-dependent exocytosis without activating a specific plant NLR receptor that guards EXO70. In this way, *Xanthomonas* efficiently inhibits the host’s PAMP-triggered immunity (PTI) by blocking exocytosis of PR1, callose deposition and the FLS2 immunity-receptor translocation to the plasma membrane, promoting successful infection.

## INTRODUCTION

Pathogens use common strategies to colonize their plant and animal hosts. Pathogenicity, for the majority of the Gram-negative bacterial species, largely depends on the highly conserved <u>T</u>ype-<u>III</u> protein <u>S</u>ecretion <u>S</u>ystem (T3SS) that delivers virulence proteins, known as “effectors”, into eukaryotic host cells. The type-III effector proteins (T3EPs) interfere with various cellular responses to the pathogen’s benefit including the inhibition of the host’s defence mechanisms (Dangl and Jones, 2019; Duxbury et al., 2016; Jo et al., 2019; Mermigka and Sarris, 2019; Staskawicz et al., 2001; Tampakaki et al., 2010). Several studies have revealed functional similarities among T3EPs from plant and animal pathogenic bacteria, suggesting that they exert similar functions in eukaryotic host cells (Büttner and Bonas, 2003; Jo et al., 2019; Staskawicz et al., 2001).

Plant pathogenic bacteria have evolved multiple T3EPs enabling evasion of recognition by the host’s immunity system, that often appear to be functionally redundant. However, the role(s) and the subcellular targets of these T3EPs remain an important open question in molecular host-microbe interactions and have only partially been elucidated.

Bacterial species of *Xanthomonas* genus are important model species for studies on the bacterial pathogenicity and host adaptation (An et al., 2020; Niño-Liu et al., 2006; Timilsina et al., 2020). This genus consists of a large group of plant-associated Gram-negative bacteria, with a high impact on food production due to severe diseases that 27 *Xanthomonas* species cause on almost 400 monocot and dicot plants (Ryan et al., 2011). *Xanthomonas campestris* pv. *campestris* infects through the vascular system the economically important plant family *Brassicaceae*, which includes the model species *Arabidopsis thaliana* (Fargier et al., 2011). The virulence of most *Xanthomonas* species depends on the presence of the *hrp* gene cluster that encodes the type-III secretion system (T3SS) and a number of T3EPs. Different *Xanthomonas* species or even different strains of the same species reveal divergent numbers of T3EPs. However, most of the sequenced species comprise of a core set of nine effectors (Ryan et al., 2011; Vicente and Holub, 2013). In *Xanthomonas campestris* the core type-III effectorome is limited to only three of these effector proteins, the XopP, XopF1 and XopAL1, indicated as Xops (*<u>X</u>anthomonas <u>o</u>uter <u>p</u>rotein<u>s</u>*) (Roux et al., 2015).

XopP was initially identified in *Xanthomonas campestris* pv. *vesicatoria* using the AvrBs2/Bs2 reporter system (Roden et al., 2004). Though of unknown function, it has being implicated in virulence in both monocot and dicot species. A XopP homolog from *Xanthomonas oryzae* pv. *oryzae* (*Xoo*) strongly suppresses peptidoglycan- and chitin-triggered immunity in rice, targeting a positive regulator of plant immunity, a ubiquitin E3 ligase, inhibiting its ligase activity and therefore suppressing *Xoo* resistance (Ishikawa et al., 2014).

When plants are exposed to pathogenic microbes, different layers of immune responses are activated. Pattern recognition receptors (PRRs) in the plasma membrane recognize conserved domains of pathogens, called pathogen-associated molecular patterns (PAMPs) and activate pattern-triggered immunity (PTI). This includes the activation of signaling cascades that leads to secretion of defense components and callose deposition for cell wall fortification. However, adapted pathogens translocate T3EPs inside the host cell that can inhibit PTI, leading to effector-triggered susceptibility (ETS) (Dangl and Jones, 2019; Duxbury et al., 2016; Jones and Dangl, 2006; Mermigka and Sarris, 2019). In turn, plants have employed intracellular nucleotide-binding domain and leucine-rich repeat (NLR) receptors to detect the effects of specific pathogen T3EPs. Protein domains that function as integrated decoys (IDs), are present in a number of NLRs. These IDs are targeted by the effectors and subsequently lead to initiation of effector-triggered immunity (ETI) (Duxbury et al., 2016; Sarris et al., 2015, 2016).

The exocyst complex is a mediator in plant immunity, tethering the post-Golgi secretory vesicles to the plasma membrane in the early stage of exocytosis, before SNAREs drive fusion of vesicles with membranes (Elias et al., 2003). Consistent with this, components of the complex are targeted by various pathogens. In plants, as in animals and other eukaryotes, the exocyst complex consists of eight subunits; SEC3, SEC5, SEC6, SEC8, SEC10, SEC15, EXO70 and EXO84. However, unlike in mammals, in plants the EXO70 subunit has 23 paralogs (Elias et al., 2003; Zhang et al., 2010). Exocyst has been reported to play an important role in PTI induction and resistance against bacteria, oomycetes and fungi in *Arabidopsis thaliana* (Peenková et al., 2011; Stegmann et al., 2012, 2014). For *Arabidopsis* EXO70B1, it has been shown to be guarded by the NLR TN2, while *exo70B1* mutants activate TN2-mediated ETI-like immunity (Zhao et al., 2015). Furthermore, interaction between *At*EXO70B1 and RIN4, a known regulator in defense responses has been reported. The latter is cleaved by AvrRpt2, an effector of *Pseudomonas syringae* and it was hypothesized that this cleavage released both RIN4 and EXO70B1 from plasma membrane to the cytoplasm, inhibiting vesicle tethering (Sabol et al., 2017). Moreover, it has been reported that AvrRpm1, from *Pseudomonas syringae*, phosphorylates RIN4 which subsequently inhibits its interaction to EXO70s, including EXO70B1, leading finally to inhibition of various defense molecules’ secretion (Redditt et al., 2019).

Here, we report that the conserved T3EP XopP from *Xanthomonas campestris* pv. *campestris* associates with various members of the exocyst complex, including EXO70B1, SEC10 and EXO84B, resulting in inhibition of the exocytosis process and consequently the host's PTI response. We showed that XopP inhibits secretion of the Pathogenesis-related protein 1A (PR1a) in transient expression assays, as well as, the callose deposition in *Arabidopsis* transgenic lines expressing XopP. Likewise, the XopP/EXO70B1 and EXO70B2 interaction affects the EXO70-mediated trafficking of FLS2 to the plasma membrane, however, without activating the TN2 dependent immunity.

This is the first report of a *Xanthomonas* core effector that targets members of the exocyst complex, potentially inhibiting its assembly. Our results support that the XopP/*At*EXO70 interaction potentially results in steric hindrance, rather than an enzymatic effect, as the molecular mechanism by which the XopP effector anchors to the exocytosis complex and manipulates exocytosis avoids activation of host NLR-dependent immunity.

## Results and Discussion

### XopP targets EXO70B1 *in vitro* and *in vivo*

NLR-IDs may reveal host proteins targeted by pathogens (Sarris et al., 2016). To further test this hypothesis, we performed a yeast two hybrid (Y2H) screen between conserved IDs from a wide range of NLRs, to detect potential interactors with the *Xanthomonas* core effectorome. We cloned twenty-four conserved plant NLR-IDs into pGBKT7-RFP vector and thirteen effectors, including nine core effectors from *Xanthomonas* species, into pGADT7-RFP vector and conducted a Y2H screen. This revealed several interactions, including an EXO70 interaction with XopP, which we decided to investigate in depth (Fig. 1A, S1D).

**Figure 1.**
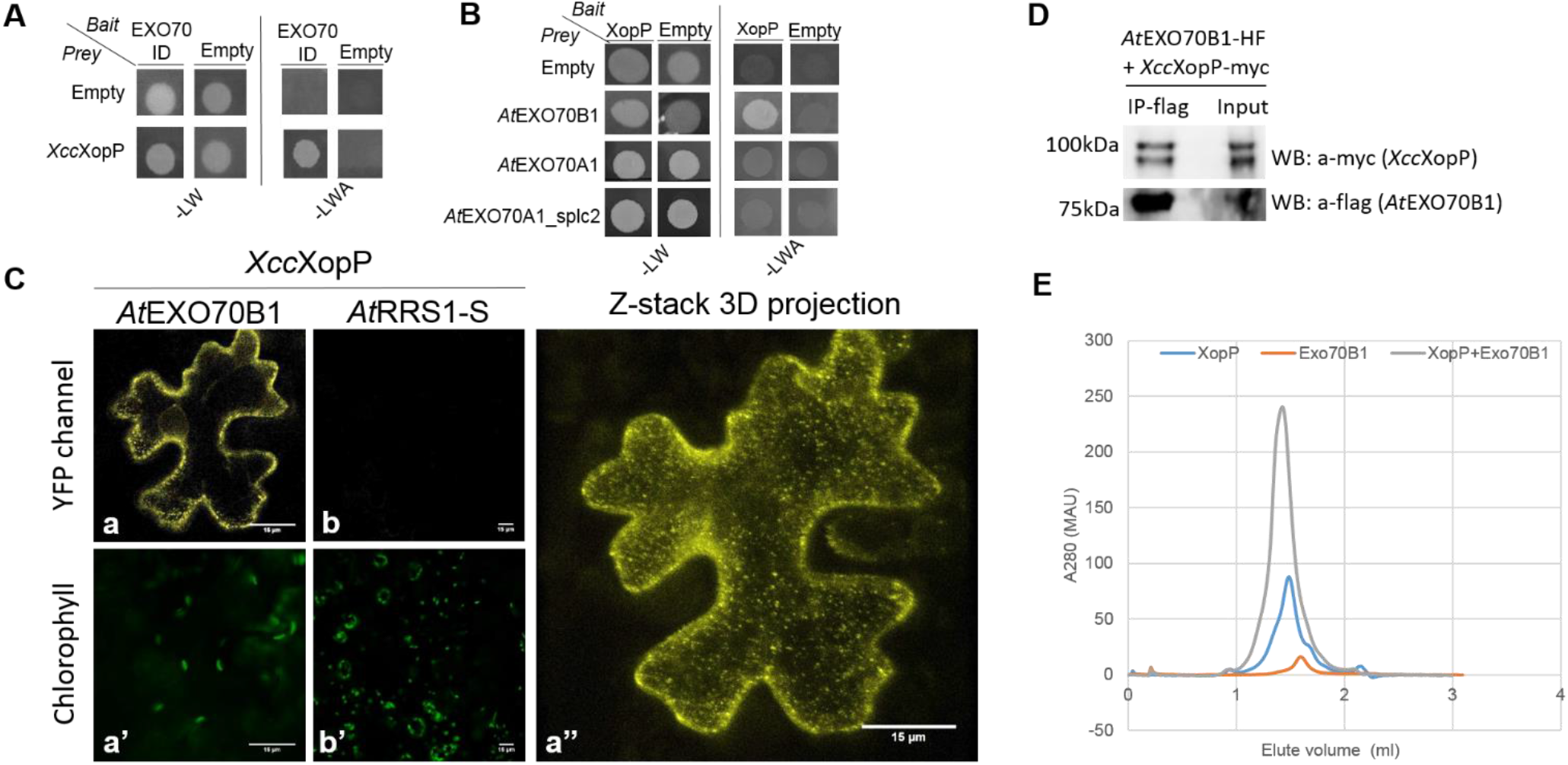
XopP targets EXO70B1 *in vitro* and *in vivo*. **A.** *Xcc*XopP targets an EXO70 integrated domain (ID) in a Y2H screen. **B.** Interaction of *At*EXO70B1 with *Xcc*XopP effector in Y2H screen. All yeast combinations grow in SC-LW medium and only *At*EXO70B1 with the effector survives in SC-LWA, in contrast to *At*EXO70A1 (both spliceforms). **C.** In planta validation of *At*EXO70B1/*Xcc*XopP interaction using a BiFC assay, in *N. benthamiana* plants. EXO70B1 and *Xcc*XopP were fused C-terminally with nVENUS and cCFP epitope tags respectively and YFP signal was observed in confocal microscopy 3dpi (a, a’’). A 3D projection of Z-stack of the interaction is shown in a’’. As negative controls, another plant protein (*At*RRS1-S) was used with *Xcc*XopP (b, b’) and no YFP signal was observed. Bars= 15μm. **D.** In planta validation of *At*EXO70B1/*Xcc*XopP interaction in *N. benthamiana* plants using a co-immunoprecipitation (co-IP) assay. EXO70B1 and *Xcc*XopP were fused C-terminally with HF (6xHis and 3xFLAG) and -myc respectively and an IP-flag was performed. *Xcc*XopP interacts with EXO70B1 as shown in the western-blot with anti-myc. **E.** SEC elution profiles for *Xcc*XopP (blue), Exo70B1 (orange) and XopP/EXO70B1 complex (grey).

XopP is a highly conserved effector (Roux et al., 2015; Ryan et al., 2011; Vicente and Holub, 2013). In general, the activities of T3EPs and their roles in virulence can be masked by redundancy. Nevertheless, a significant reduction in virulence was previously reported for a XopP Tn5gusA5 insertion mutant of the model strain *Xanthomonas campestris* pv. *campestris* 8004. The mutated strain reveals significantly reduced symptoms of black rot on the leaves of Chinese radish (*Raphanus sativus* var. *radiculus*) compared to the wild-type strain *Xcc* 8004 (Jiang et al., 2009). We repeated the infection assays to Chinese radish plants using the wild-type strain *Xcc* 8004 and the XopP mutant and we obtained similar results (Fig. S5C).

To determine whether EXO70 is a real target of XopP, we performed a BLASTP analysis using the EXO70-ID as a query. From both protein and nucleotide blast analysis, most hits were EXO70B1. Thus, we cloned EXO70B1 from *Arabidopsis thaliana* in pGBKT7 vector and performed a yeast two hybrid assay with the effector. As shown in Fig. 1B and S1E, *At*EXO70B1, but not *At*EXO70A1 (splice forms 1 and 2), strongly interacts with XopP, indicating that this could be an authentic target. Our results also indicate a specificity to EXO70B1. To further reinforce this observation, we investigated the co-localization of the two proteins (Sabol et al., 2017). We constructed XopP-mCherry and *At*EXO70B1-mCherry fusions at the C-terminus and transiently expressed the constructs under the 35S CamV promoter in *Nicotiana benthamiana* leaves. We observed the same compartmentalization of both XopP and *At*EXO70B1 in the cell cytoplasm and inside the nucleus (Fig. S1A).

To validate the interaction in planta, we performed Biomolecular Fluorescence Complementation (BiFC) and co-immunoprecipitation (co-IP) analyses (Fig 1C and 1D, respectively). For BiFC assay, we transiently expressed under the CaMV 35S promoter in *Nicotiana benthamiana* leaves the EXO70B1 or the NLR RRS1-S (used as a negative control) fused with nVENUS C-terminally, together with the XopP or the AvrRps4 effector from *Pseudomonas syringae* pv. *pisi* (as a negative control), both fused with cCFP C-terminally. EXO70B1 interacted specifically with XopP (Fig. 1Ca) but not with AvrRps4 (Fig. S1B) and the complex is localized to the cytoplasm, but also in the inner periphery of the plasma membrane, as it is shown by spots in 3D projection (Fig 1Ca’’). XopP did not interact with RRS1-S (Fig. 1Cb). For the co-IP assay, we transiently expressed EXO70B1 fused to a tandem 6xHis and 3xFLAG epitope tag (HF) and XopP fused with a -Myc tag and performed co-IP with a-FLAG beads. In Fig. 1D, the interaction of these two proteins is observed, while as a negative control the AvrRps4 effector was used (Fig. S1C, upper and lower lanes, respectively).

To investigate if the two proteins interact also *in vitro*, size exclusion chromatography (SEC) experiment was performed to XopP and EX070B1 proteins separately and together, using the same type of column (Superdex™ 200 Increase 3.2/300). As shown in Fig. 1E, when the protein solutions were combined at a ratio of 1:1 both proteins were eluted in earlier fraction volumes, in comparison to their own elution profiles. This strongly suggests an *in vitro* complex formation between XopP and EXO70B1.

Furthermore, we investigated the potential co-localization of the XopP itself, as well as the *Xcc*XopP/*At*EXO70B1 BiFC complex, with the autophagosome marker ATG8-mCherry, a <u>P</u>lasma <u>M</u>embrane (PM) marker and the early/late endosomal marker, *Ds*Red-FYVE, using confocal microscopy. However, we did not manage to show these proteins to be co-localized with any of the above markers, except for PM-marker (Fig. S7A, B).

### EXO70B1 specifically interacts with XopP and not with other homologs

The next step was to determine whether EXO70B1 interacts with other XopP homologs or if this interaction is limited specifically to XopP originating from *Xanthomonas campestris* pv. *campestris* (here after *Xcc*XopP). We initially performed a BLASTP search at the NCBI and KEGG in order to identify the *Xcc*XopP homologs. Next we performed a phylogenetic analysis of all XopP homologs, using a Maximum-Likelihood method. The analysis was not limited to *Xanthomonas* species, since we identified homologs to *Ralstonia* and *Acidovorax* species. Our results pointed out five subgroups, including the *Ralstonia* group, the *Acidovorax* group and three *Xanthomonas* groups indicated as XopP1, XopP2 and the EXO70 interacting *Xcc*XopP groups (Fig. 2A). We amplified from each of the four groups a representative effector to test in a yeast-two hybrid analysis whether they interact with EXO70B1. We used two effectors from previous work of our lab, the *Xanthomonas oryzae* pv. *oryzicola* XopP1 and XopP2 (Michalopoulou et al., 2018) and *Ralstonia solanacearum* Hlk3 and *Acidovorax citrulli* XopP-homologs. The tested preys are represented in the phylogenetic tree in bold (Fig. 2A). In our analysis, we also included *At*EXO70F1. Interestingly, EXO70B1 did not interact with any homologs of XopP used, whereas EXO70F1 interacted with *Xoc*XopP1 (Fig. 2B), which indicates a specificity regarding XopP/ EXO70 interactions.

**Figure 2.**
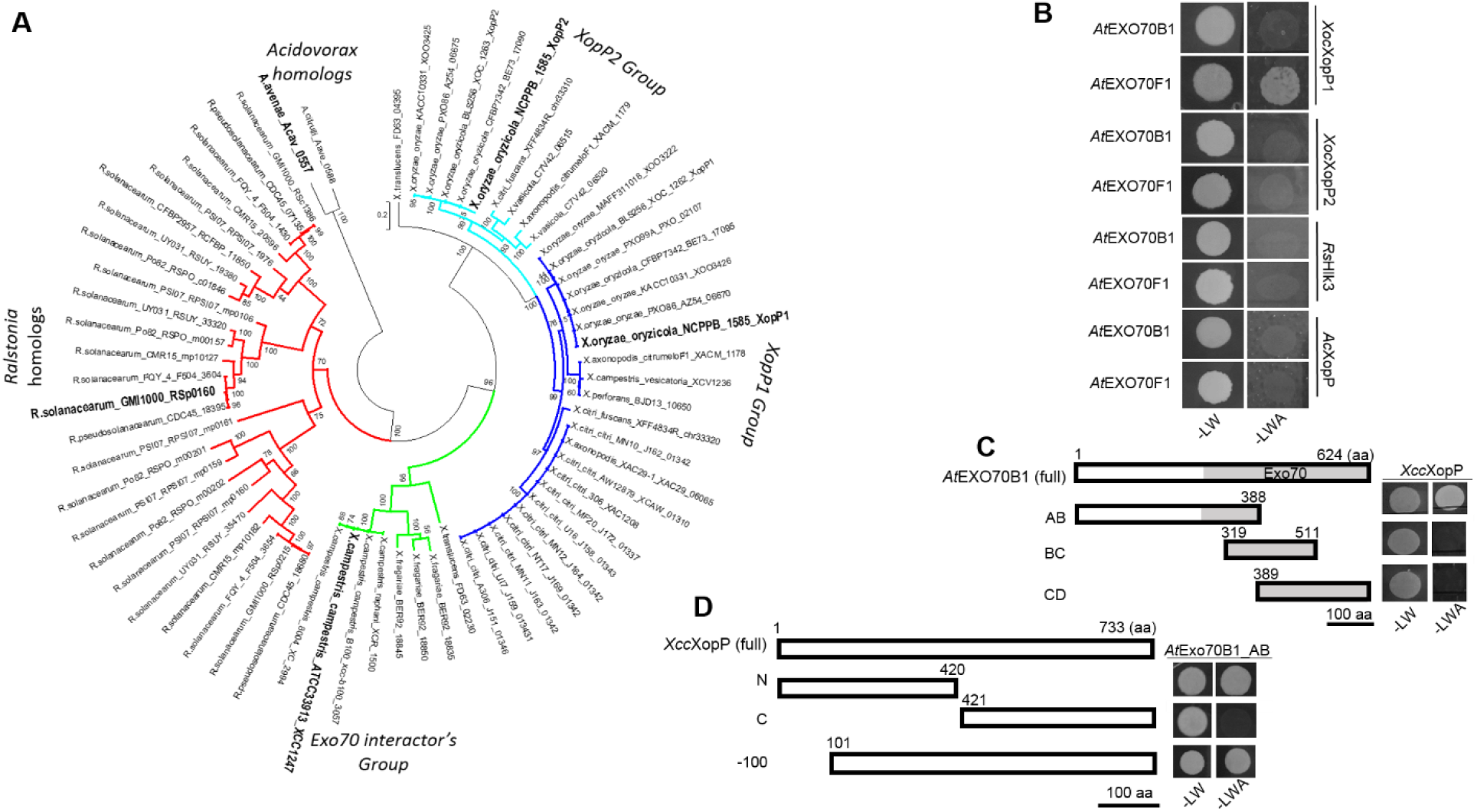
EXO70B1 interacts only with XopP from *Xanthomonas campestris* pv. *campestris* and not with its orthologs and the interaction is through their N-terminal parts. **A.** Phylogenetic analysis of XopP homologs. XopP homologs are grouped in different clusters. The homologs presented in bold were used for the interaction screening. **B.** EXO70B1 interacts specifically with *Xcc*XopP and not with other orthologs. In contrast, EXO70F1 interacts only with *Xoc*XopP1. **C.** Schematic representations of *At*Exo70B1 and XopP truncations. With grey the catalytic domain (265 to 614 aa) of Exo70 is presented. Bar = 100 aa. Next to each construct appear the results of the Y2H assays. Only *At*EXO70B1-AB truncation, which corresponds to the first 388 aa, interacts with full-length *Xcc*XopP (upper panel) as it is observed in SC-LWA medium. **D.** *Xcc*XopP N-terminal domain interacts with *At*EXO70B1-AB truncation. Similarly, the full-length *At*EXO70B1 interacts with *Xcc*XopP truncation lacking the first 100 aa (XccXopP-100), as it is observed in SC-LWA medium.

### EXO70B1 and XopP interact at their N-termini

We sought to further characterize the interaction between EXO70B1 and *Xcc*XopP, in order to minimize the area of binding capacity. For this, we constructed three truncations of the plant protein according to domains similarly as for *At*EXO70B2 (Fig. 2C) (Teh O.K. et al., 2019). The truncated proteins were expressed in the yeast vector as baits and tested for interaction with the effector via Y2H. Only AB domain, which corresponds to the first 388 amino acids of the protein, interacts with the effector (Fig.2C), while the expression of the other constructs was verified via western blot analysis (Fig. S3C). We next split the *Xcc*XopP in N- and C-terminal fragments (Fig. 2D), using Phyre2 bioinformatics program (Kelley et al., 2016), as there is no information of the effector’s structure. Similarly, Y2H showed that EXO70B1 and the effector interact at their N-terminal parts (Fig. 2D). In addition, during the co-IP analysis we observed that two cleaved forms of XopP are present (Fig. S2A). Based on this observation we decided to clone *Xcc*XopP minus the first 100 amino acids, leading to construct *Xcc*XopP-100 where cleaved forms no longer appear (Fig. 2D and S2A). This is also consistent with previous reports that many T3EPs are cleaved in their first amino acids when entering host cells, in order to be activated (Mudgett et al., 2000; Sohn et al., 2009). Next, we checked that *Xcc*XopP-100 continues to interact with EXO70B1 in Y2H assay (Fig. 2D). The *Xcc*XopP-100 appears to be uncleaved, and interestingly, in visualization of field-emission SEM (FE-SEM) of a gel-like formation of *Xcc*XopP-100 after dialysis in storage buffer in absence of β-mercaptoethanol, we observed the formation of fibrils (Fig. S2B). Similar fibrils were observed in TEM micrographs of *Xcc*XopP-100 after overnight dialysis in storage buffer in absence of β-mercaptoethanol (Fig. S2C).

In yeast and mammals, it has been shown that EXO70B1 interacts with phosphatidylinositol 4,5-bisphosphate (PI(4,5)P2) of the plasma membrane (PM) through its basic residues, localized at C-terminus (Wu and Guo, 2015). Recently, this association with the PM has been extended also for plants. Endosidin2 (ES2), a small molecule, inhibited the EXO70A1 binding to the PM (Zhang et al., 2016). From these data and according to our results, we conclude that XopP does not target the C-terminal part of EXO70B1 to directly inhibit this association although we cannot exclude that the effector’s binding at the N-terminus of EXO70B1 might change its whole conformation.

### *Xcc*XopP associates with *Arabidopsis* EXO70F1 and EXO70B2 but not with the Exo70B1 from monocots

EXO70B1 is a conserved part of the exocyst complex, being implicated in immune responses to different pathogens and autophagy (Kulich et al., 2013; Stegmann et al., 2014; Zhao et al., 2015). In an attempt to investigate if the *Xcc*XopP also targets other EXO70s that are also involved in immune responses (Peenková et al., 2011; Stegmann et al., 2012), we chose two paralogs of *exo70B1*, the phylogenetically close *At*Exo70B2 and the phylogenetically distant *At*EXO70F1 for Y2H analyses (Stegmann et al., 2012). We have already seen that *At*EXO70F1 interacted with *Xoc*XopP1 (Fig. 2B). Moreover, we checked the intraspecies specificity of the effector; and we cloned *Exo70B1* from the monocot *Oryza brachyantha,* a wild-crop relative of common rice. Interestingly, during the cloning process, we discovered a smaller, not annotated version of the protein, lacking the first 234 amino acids. We cloned this short version of *Obexo70B1* from cDNA (termed *Ob*EXO70B1_small). Notably, the large version of the protein was not expressed in our conditions and needed to be cloned using gDNA (termed *Ob*EXO70B1_large) (Fig. S3A). All EXO70B1 proteins were expressed as baits. In a Y2H screen we observed that both *At*EXO70B2 and *At*EXO70F1 interact with *Xcc*XopP. However, both *Ob*EXO70B1_small and *Ob*EXO70B1_large do not interact with the effector (Fig. S4A). *Ob*EXO70B1_small and large protein expression was verified via western blot analysis (Fig. S3B). This finding led us to perform an amino acid alignment between all *Xcc*XopP interacting and non-interacting EXO70s, including the NLR-ID, in the hope of identifying possible amino acid changes in the non-interacting protein. We focused on the N-terminal domain of *At*EXO70B1 which interacts with *Xcc*XopP. For the alignment we used only the *Ob*EXO70B1_small protein sequence. Even if the N-terminal part of the proteins seems to be quite divergent, we identified one crucial amino acid change, from lysine to glutamate, that seems to be conserved among the *Xcc*XopP-interacting homologs but appears to be mutated in *Ob*EXO70B1_small (Fig. S4B).

To test if glutamate is an essential amino acid for the *Xcc*XopP/EXO70 interaction, we constructed mutated forms of *Ob*EXO70B1_small and *At*EXO70B1, corresponding to K134E and E347K, respectively and tested again for interaction with *Xcc*XopP in a two-hybrid analysis in *Saccharomyces cerevisiae* (Fig. S4C). Interestingly, we observed a partial gain of interaction between the mutated *Ob*EXO70B1_small^K134E^ and the effector. However, the mutated *At*EXO70B1^E347K^ still interacted with *Xcc*XopP. Thus, glutamate seems to be a necessary but not sufficient amino acid for *Xcc*XopP/EXO70 interaction, since its mutation does not abolish the interaction. Future structural studies will reveal the exact nature of this interaction.

In plants, the EXO70 subunit of the exocyst complex comprises a large number of paralogs. For instance, in *Arabidopsis thaliana* there are 23 different EXO70 isoforms. This led to the hypothesis that different EXO70 subunits have specialized functions inside the cells and may take part of specific cargoed-vesicle exocytosis (Saeed et al., 2019; Sekereš et al., 2017) The fact that XopP associates with EXO70s from different phylogenetic clades could indicate that this core effector of *Xanthomonas* species might target the exocytosis of different cargos, such as antimicrobial compounds, callose or other membrane and lipid features.

### *Xcc*XopP inhibits the exocytosis of <u>P</u>athogenesis-<u>R</u>elated protein <u>1A</u> (PR1a)

The exocyst complex mediates the tethering of secretory vesicles to the cell plasma membrane (Guo et al., 1999). Under pathogen attack, this complex also contributes to efficient immunity by sending the TGN-originating secretory vesicles or MVBs originating from the process of autophagy, to the site of the attack (Pečenková et al., 2017). Furthermore, it has been reported that the exocyst is required for secretion of the Pathogenesis-Related protein 1A (PR1a) (Du et al., 2015; Gu and Innes, 2012; Hammond-Kosack et al., 1994). Our finding that *Xcc*XopP targets *At*EXO70B1, *At*EXO70B2 and *At*EXO70F1 (Fig. 1B; S4A), led us to speculate that the effector may act as an inhibitor of exocytosis during immune activation. To check our hypothesis we initially investigated if the PR1a secretion is compromised by *Xcc*XopP. For this we used the Chp7 serine protease effector from *Clavibacter michiganensis subsp. sepedonicus* that has been shown to induce an apoplastic Hypersensitive Response (HR) in *Nicotiana tabacum* only when it is targeted to the apoplast via the PR1a secretion peptide, but not intracellularly (Lu et al., 2015). We transiently co-expressed the PR1sp-Chp7 construct together with *Xcc*XopP effector in *Nicotiana tabacum* leaves. Both constructs were expressed using the 35S CaMV promoter. As a negative control we used the GUS protein. Three days post infiltration (dpi), an HR was observed in the leaves where the PR1sp-Chp7 was co-expressed with GUS, but not along with *Xcc*XopP (Fig. 3A). To rule out any possibility of *Xcc*XopP being mediated as a general cytoplasmic HR inhibitor we performed a co-expression of *Xcc*XopP with the *Xcv*XopQ effector that induces cytoplasmic HR in *Nicotiana tabacum* leaves (Adlung et al., 2016). Similarly, to rule out any potential interaction of *Xcc*XopP with Chp7, we performed a Y2H analysis between *Xcc*XopP as a bait and the Chp7 as a prey. As it is shown in Fig S5A and B, the *Xcc*XopP did not inhibit the *Xcv*XopQ-dependent cytoplasmic HR, nor associated with Chp7 in a Y2H assay, respectively. Overall, these results indicate that *Xcc*XopP inhibits the PR1a exocytosis process, probably via interacting with EXO70B1 or any other(s) member(s) of the exocytosis complex.

**Figure 3.**
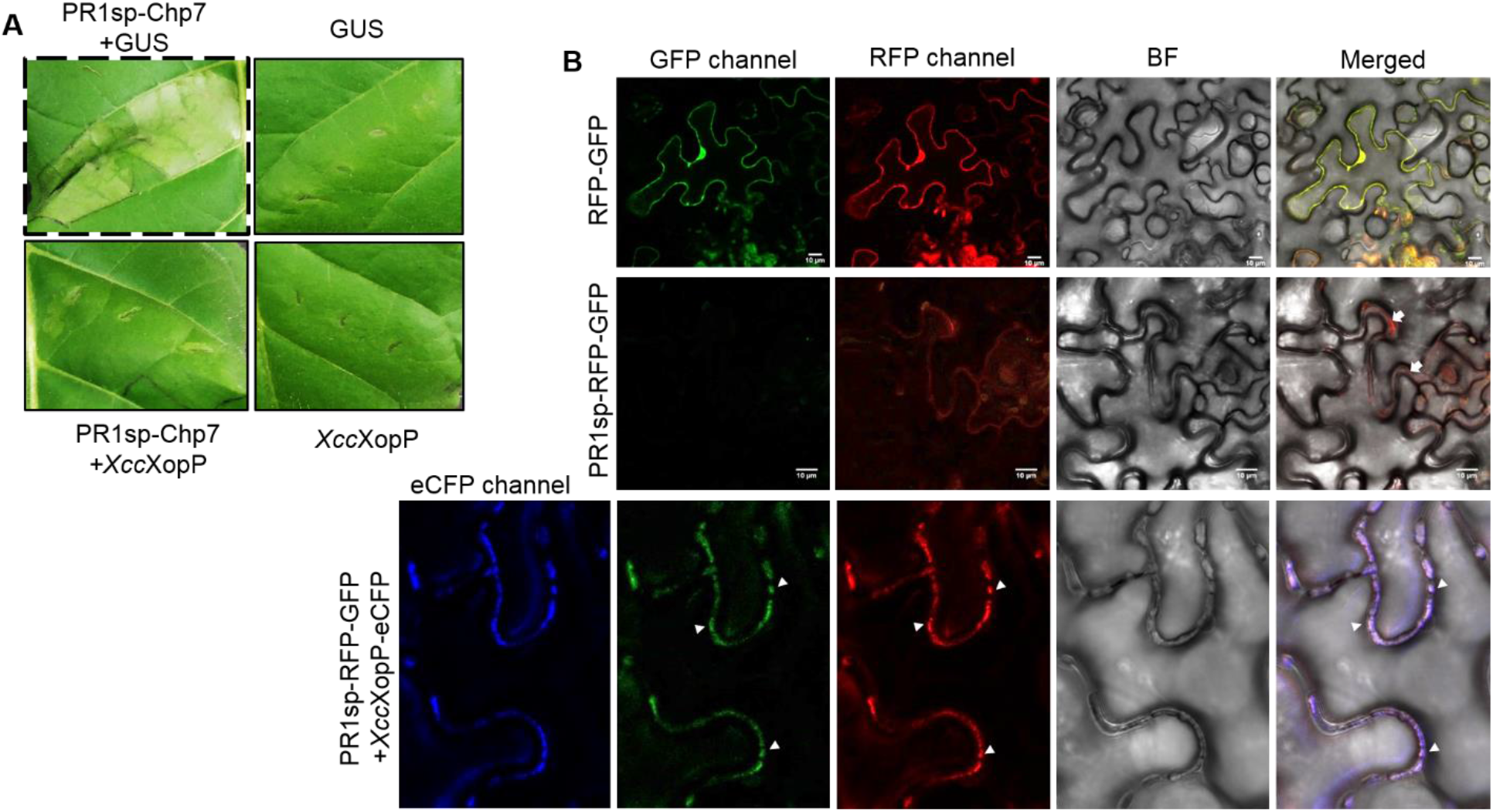
*Xcc*XopP inhibits the exocytosis of Pathogenesis-Related protein 1A (PR1A). **A.** HR inhibition assay. *Xcc*XopP inhibits the exocytosis of the chimeric protein PR1a-Chp7, to the apoplast. Inhibition of HR (left lower lane) appears 3dpi in *N. tabacum* leaves. GUS protein was used as an HR negative control. Expression of GUS or XopP itself does not produce any HR (right lanes). The experiment has been repeated three times with identical results (total plants used in which HR was inhibited = 11/11). **B.** The chimeric protein RFP-GFP is expressed in the cytoplasm (upper panel). The PR1a secretion peptide fusion to RFP-GFP chimera (PR1sp-RFP-GFP) results in exocytosis to the apoplast of the RFP-GFP (white arrows, middle panel). Only RFP is visualized, as GFP is quenched in the acidic environment of the apoplast. When *Xcc*XopP is expressed (eCFP channel) along with PR1sp-RFP-GFP, the RFP-GFP exocytosis is inhibited and it is relocalized to the cytoplasm, as both fluorescence proteins are visible and localized in vesicle-like formations (white arrowheads). The pictures are taken in a confocal microscope, 3dpi of *N. benthamiana* leaves. Bars = 10 μm. The experiment was repeated three times with similar results.

We used confocal microscopy to further test if *Xcc*XopP inhibits PR1a exocytosis. After N-terminally fusing the PR1 secretion peptide with the chimeric RFP-GFP protein (35S::PR1sp-RFP-GFP), we transiently co-expressed the 35S::PR1sp-RFP-GFP in *Nicotiana benthamiana* leaves with and without *Xcc*XopP effector using the GUS protein as a negative control. When the RFP-GFP chimeric protein lacking a cell secretion peptide is expressed, it reveals a nuclear-cytoplasmic localization, so, both RFP and GFP signals are detected (Fig. 4B). Contrariwise, when the chimeric RFP-GFP protein was targeted to the apoplast via the PR1sp, we predominantly observed RFP signal, since the GFP is almost quenched in the acidic environment of the apoplast (Fig. 4B). When *Xcc*XopP was co-expressed with PR1sp-RFP-GFP both fluorescent signals were co-localized near the plasma membrane in vesicle-like formations, indicating the inhibition of the chimeric PR1sp-RFP-GFP translocation to the apoplast (Fig. 4B; Fig. S5F).

**Figure 4.**
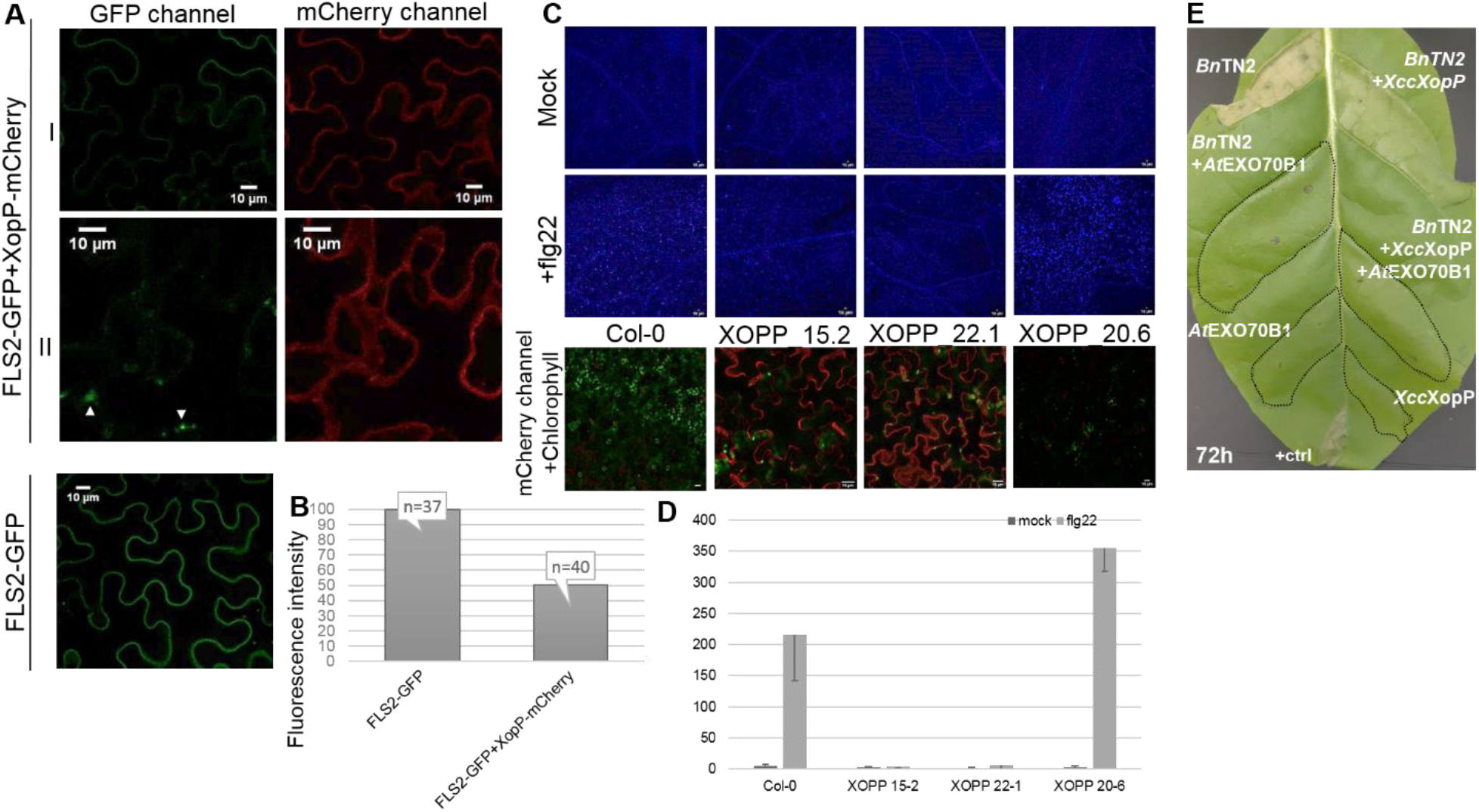
XopP inhibits the membrane translocation of the transmembrane immunity receptor FLS2 and Callose deposition, without activating TN2-mediated immunity. **A, B.** Visualization (A) and quantification (B) of representative pictures of expression of FLS2 receptor fused C-terminally to GFP tag in the plasma membrane. The FLS2-GFP is co-expressed with *Xcc*XopP fused to mCherry tag (left) or expressed alone (right). In samples of subgroup-I, half of the FLS2 membrane localization is observed, while in samples of subgroup-II almost no FLS2 is visible in the plasma membrane, while specific foci are present, resemble vesicle-like formations that cannot be exocytosed (white arrowheads). The pictures are taken in a confocal microscope 3dpi in *N. benthamiana* leaves and are representatives from total 40 cells. Bars = 10 μm. The intensity measurement has been made with Image J software and the number of cells appear above its column. **C, D.** Visualization (C) and quantification (D) of callose deposition after elicitation with 20μm flg22 (middle lane) or mock solution (upper lane), in T1 lines of transgenic *Arabidopsis thaliana* (15-2, 22-1 and 20-6) overexpressing *Xcc*XOPP. In the lower lane, *Xcc*XopP expression is shown in confocal microscope and it is in accordance with the levels of Callose secretion. In line 20.6, the *Xcc*XopP expression is almost absent. Bars = 15μm. The quantification has been made with Image J software and data represent the mean +/− SD. **E.** Transient expression of the NLR immune receptors *Bn*TN2 and *At*EXO70B1, and *Xcc*XopP in *N. tabacum leaves*. *Bn*TN2 triggers an HR, which is suppressed by *At*EXO70B1. *Bn*TN2 co-expressed with *Xcc*XopP also induces an HR reaction, while the combination of all three constructs when co-expressed, no HR reaction is observed after 72hpi. The experiment has been repeated three times with identical results.

### *Xcc*XopP inhibits the membrane translocation of the transmembrane immunity receptor FLS2

It has been recently reported that both the *Arabidopsis thaliana* EXO70B1 and EXO70B2 regulate the trafficking of the transmembrane immunity receptor FLS2 to the plasma membrane (PM) (Wang et al., 2020). FLS2 receptor is responsible for the recognition of the bacterial flagellin (flg22) epitope and the activation of the flagellin-dependent PTI in Arabidopsis (Wang et al., 2020). We wanted to determine whether *Xcc*XopP acts as a negative regulator of the EXO70B-dependent FLS2 translocation to the PM, leading to reduced FLS2 accumulation at the PM. To investigate this, we analyzed the fluorescence intensity of the chimeric FLS2-GFP protein itself or when co-expressed with *Xcc*XopP-mCherry, in *N. benthamiana* plants. When *Xcc*XopP is present, we observed a significant reduction of the FLS2 signaling in PM (Fig. 4A, B). We categorized our results in two subgroups (Fig. 4A). In subgroup I (the most abundant subgroup) and subgroup II, we observed a reduction in FLS2 signaling about 50% and 100%, respectively.

### *Xcc*XopP targets the EXO70B avoiding activation of its guarding NLR immune receptor

*Xcc*XopP is the first *Xanthomonas* effector shown to target an exocyst component to inhibit the exocytosis process, although it has been previously shown for AVR1 effector, from *Phytopthora infestans* to interact with the exocyst component Sec5 disturbing vesicle trafficking (Du et al., 2015). Similarly, the AvrPii, a fungal effector from *Magnaporthe oryzae* has been reported to interact with EXO70F2 and F3 of rice (Fujisaki et al., 2015). Recently, it was reported that the EXO70B1 is guarded in *Arabidopsis* by the NLR immune receptor TN2. The ubiquitination of EXO70B1 by the AvrPtoB effector from *Pseudomonas syringae*, leads to a TN2 activation and HR induction in *Arabidopsis* and transiently in *N. tabacum* (Wang et al., 2019).

In order to investigate if the *Xcc*XopP is able to activate TN2 immunity, we cloned the Arabidopsis TN2 resistance gene (*At*TN2-YFP) and its homolog from *Brassica napus* (*Bn*TN2-YFP). We transiently expressed in *Nicotiana tabacum* cv. Petit Gerard leaves the TN2s alone or with *At*EXO70B1. Our results revealed an HR induction by the *Bn*TN2 alone (Fig. 4E) but not by the *At*TN2 (data not shown). We verified the expression levels for both TN2 proteins using confocal microscopy (Fig. S5E). The *Bn*TN2-triggered HR was abolished when co-expressed with *At*EXO70B1 in a 1:1 ratio (Fig. 4E). Interestingly, and unlike AvrPtoB, the *Xcc*XopP did not activate the TN2-dependent HR when transiently co-expressed in *N. tabacum* (Fig. 4E). This result led us to assume that *Xcc*XopP does not damage EXO70B1 protein. This was expected since no cleaved forms of the protein appeared in the co-IP assay (Fig. 1D). We believe that *Xcc*XopP associates with the exocyst component, potentially deploying a different mechanism to inhibit exocytosis.

To further verify our observation, we generated *Arabidopsis thaliana* transgenic lines expressing the *Xcc*XopP-mCherry under the regulation of the Arabidopsis Rubisco promoter and the NOS terminator, in cultivar Col-0. Our transgenic lines did not reveal any TN2-dependent phenotype (Fig, S5D), unlike the AvrPtoB transgenics which were lethal due to TN2 activation (Wang et al., 2019).

### *Xcc*XopP inhibits cell wall callose deposition through exocyst targeting

When PAMPs are recognized by PRR receptors, cellular events such as activation of MAP kinases, production of ROS and callose deposition are activated as a regular PTI response. Callose deposition is a common target for many pathogens, including several *Xanthomonas* species, which inhibit flg22-callose deposition at the cell wall, however with an unknown mechanism for the majority of them (Popov et al., 2016). Furthermore, it has been reported that the callose secretion/deposition in plants involves the exocyst complex (Du et al., 2015; Nielsen et al., 2012; Robatzek, 2007; Xu and Mendgen, 1994).

We evaluated callose deposition in *Arabidopsis thaliana* transgenic lines that stably express *Xcc*XopP, using T1 plants of three of our transgenic lines to further study callose deposition levels (Fig. 4C,D), upon eliciting PTI with 20 μm flg-22. We compared our results with WT *Arabidopsis* Col-0 plants and noticed that in XopP-lines 15-2 and 22-1, almost no callose is secreted, while in XopP-line 20.6 the callose deposition levels are half of the respective WT Col-0 treated plants (Fig. 4C,D). Based on these results we evaluated the *Xcc*XopP-mCherry expression levels in all three lines using confocal microscopy. We observed that callose deposition inversely correlates with *Xcc*XopP-mCherry expression levels (Fig. 4C). Taken together, these data indicate that *Xcc*XopP inhibits the exocytosis process responsible for the callose deposition in our transgenic Arabidopsis.

### *Xcc*XopP inhibits the exocytosis through association with multiple components of the exocyst complex

The exocyst complex has been shown to form two sub-complexes (I and II), consisting of the components Sec3, Sec5, Sec6, Sec8 and Sec10, Sec15, EXO70, EXO84, respectively (Heider et al., 2016; Mei et al., 2018). Moreover, the assembly between the exocyst components is conserved from *Saccharomyces cerevisiae* to mouse and the two sub-complexes arrive at the PM independently in mammalian cells, being able to carry vesicles without assembling in an octameric complex (Ahmed et al., 2018). In yeast, Sec10 and Sec15 form a stable pair, which then requires EXO84 to associate with EXO70 and form the full sub-complex II. Whether this model is precisely reproducible in plants remains to be discovered. To test if *Xcc*XopP has a “larger” target such as a whole sub-complex of the exocyst, we cloned the other seven exocyst components of *Arabidopsis thaliana* (including the *Sec15A*, *Sec15B*, *Exo84B* and *Exo84C* paralogs) as preys in pGADT7 vector. We investigated, in Y2H, the interactions of *At*EXO70B1 with the other exocyst components and with itself and we found that EXO70B1 interacts with SEC5A, SEC15B, EXO84B as it was previously reported (Kulich et al., 2013; Zhao et al., 2015), as well as, with itself (Fig. S6A, C). Our results reveal that the interactions of EXO70B1 with the other three exocyst components are mediated by the first 388 amino acids of EXO70B1 (AB truncation) (Fig. S6E). We also noticed a weak interaction between EXO70B1 and SEC3A, only when we transformed the AH109 yeast strain with both plasmids but not in PJ69 yeast strain. These results are in accordance with (Zhao et al., 2015). Also, using Y2H assays we investigated the interactions between the components of sub-complex II and we found that similarly to yeast and mammals, SEC15B interacts with both SEC10 and EXO84B (Fig. S6F). Our results are consistent with a previously reported Y2H screening (Hála et al., 2008; Picco et al., 2017). Furthermore, we validated *in planta* all the above interactions, using BiFC assay (Fig. S6B, F). Notably, *Xcc*XopP seems to interact with two more components of the exocyst complex, the SEC10 and EXO84B, and in fact they belong to the same sub-complex that also includes EXO70B1 (Fig. S6A). We validated our results obtained from Y2H analysis, using *in planta* BiFC assays, where *Xcc*XopP and SEC10 appear to associate in specific foci (Fig. S6B), resembling the vesicle-like formations observed when PR1-RFP-GFP’s exocytosis is inhibited (Fig. 3B; S5F). We also investigated which protein domains of *Xcc*XopP are essential for the association with both SEC10 and EXO84B, using Y2H assays. We noticed that, similarly to EXO70B1, the N-terminal region of *Xcc*XopP is responsible for these associations (Fig. S6D). Additionally, we investigated our hypothesis that *Xcc*XopP could act as an inhibitor of the assembling of the two exocyst sub-complexes or the disassembly of sub-complex II, inhibiting EXO70B1 (SC II) and SEC5A (SC I) interaction (Mei et al., 2018) or inhibiting the association of SC II components (Mei et al., 2018), respectively. To investigate our first hypothesis, we cloned the *AtExo70B1* in MCS I and *XccXopP* in MCS II of pBridge vector and transformed the Yeast-3-Hybrid (Y3H) compatible AH109 yeast strain, along with pGADT7 expressing SEC5A and we investigated the potential inhibition in Y3H assay. When L-Methionine was absent from the yeast growing medium, *Xcc*XopP was induced by the MET25 promoter and partially inhibited the EXO70B1-SEC5A association (Fig. 5). To test our second hypothesis, we cloned the *AtSec15B* in MCS I and *XccXopP* in MCS II of pBridge vector and we performed a Y3H assay between SEC15B and its interactors EXO70B1, SEC10 and EXO84B. We observed a partial inhibition of the association between SEC15B and SEC10 (Fig. 5), while, SEC15B did not interact with EXO70B1 when transformed as bait and prey, respectively. Thus, we tested the potential inhibition generating the opposite constructs, where EXO70B1 is cloned as a bait in pBridge and SEC15B is cloned as a prey in pGADT7, vectors respectively. As we present in Fig. 5, we observed a partial inhibition by *Xcc*XopP. Furthermore, we tested the inhibition of SEC15B and EXO70B1 interaction by *Xcc*XopP, via triple co-IP analysis. We cloned *Sec15B* fused to HF C-terminal epitope tag and *Exo70B1* and *XccXopP* fused to -myc C-terminal epitope tag. We performed a triple IP with FLAG-beads and we checked the EXO70B1 levels via western blot analysis, in presence and absence of *Xcc*XopP. As it is shown in Fig. S7C, we did not succeed to visualize the inhibition effect in presence of XccXopP. However, it is noteworthy that XccXopP co-IPed with EXO70B1, although it does not directly interact with SEC15B. From this assay we cannot observe the inhibition we observed in Y3H assay. This can be explained, since co-IP does not represent a quantitative method and it is difficult to see the inhibition we observed in Y3H assays.

**Figure 5.**
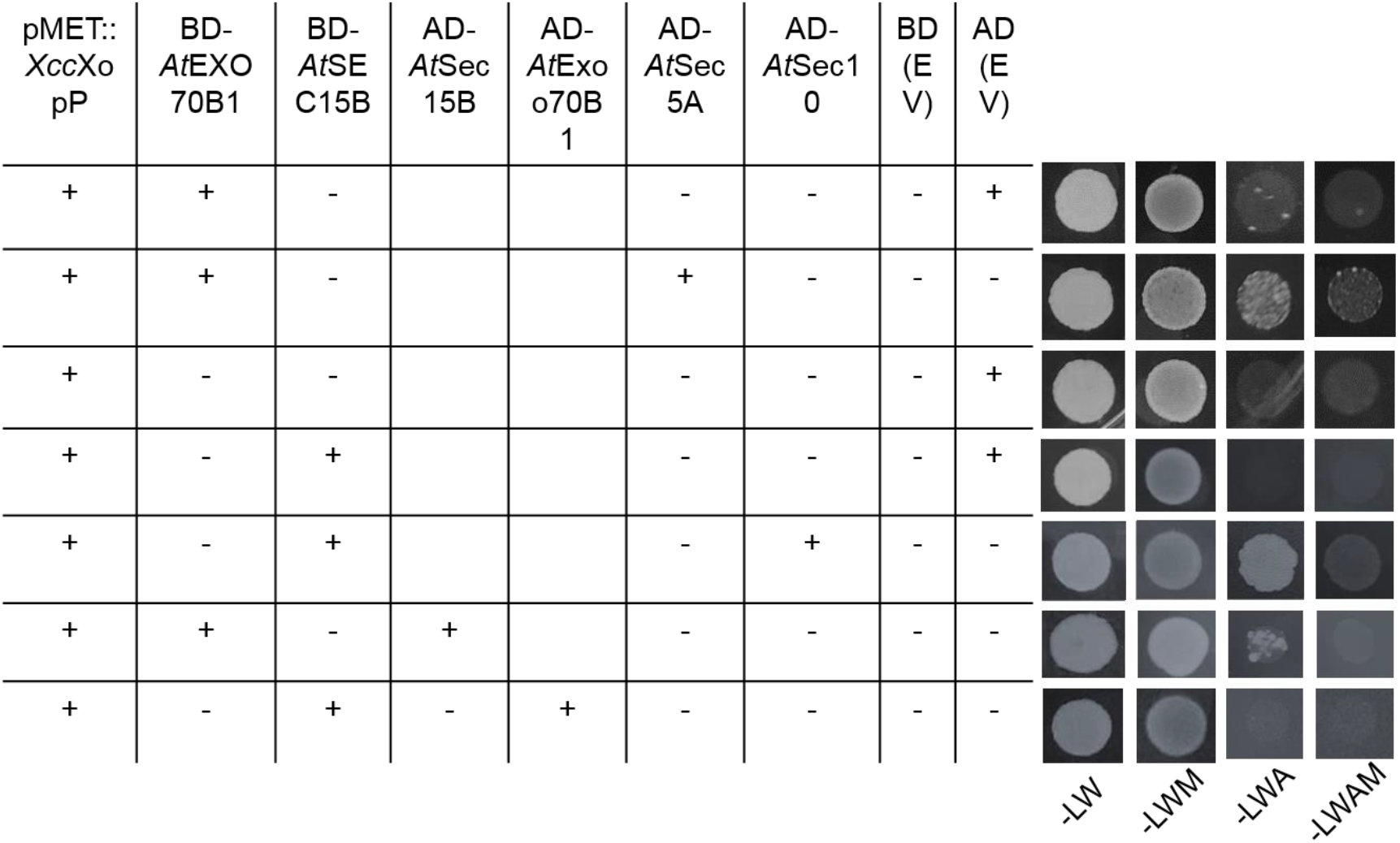
*Xcc*XopP inhibits the EXO70B1/SEC5A, SEC15B and SEC15B/SEC10 associations in yeast-three-hybrid (Y3H) assays. *Xcc*XopP interferes with EXO70B1/SEC5A, EXO70B1/SEC15B and SEC15B/SEC10 associations in a yeast-three-hybrid analysis. The constructs were transformed in the Y3H compatible yeast strain AH109. EXO70B1 associates with SEC5A and SEC15B and the latter associates with SEC10, as it is shown in medium lacking leucine, tryptophan and adenine. When methionine is absent from the medium, *Xcc*XopP is expressed by a MET25 inducible promoter and the above associations are inhibited.

Our overall data, can reasonably lead us to speculate that the effector acts as an intermediate molecule, inhibiting sub-complex II from forming a functional complex with sub-complex I. Future structural studies will examine if the effector can mimic EXO70B1 and if they share a similar structure and dimerize (Fig.S6A, C), enabling *Xcc*XopP to replace it with a non-functional component at the sub-complex II (Fig. 6).

**Figure 6.**
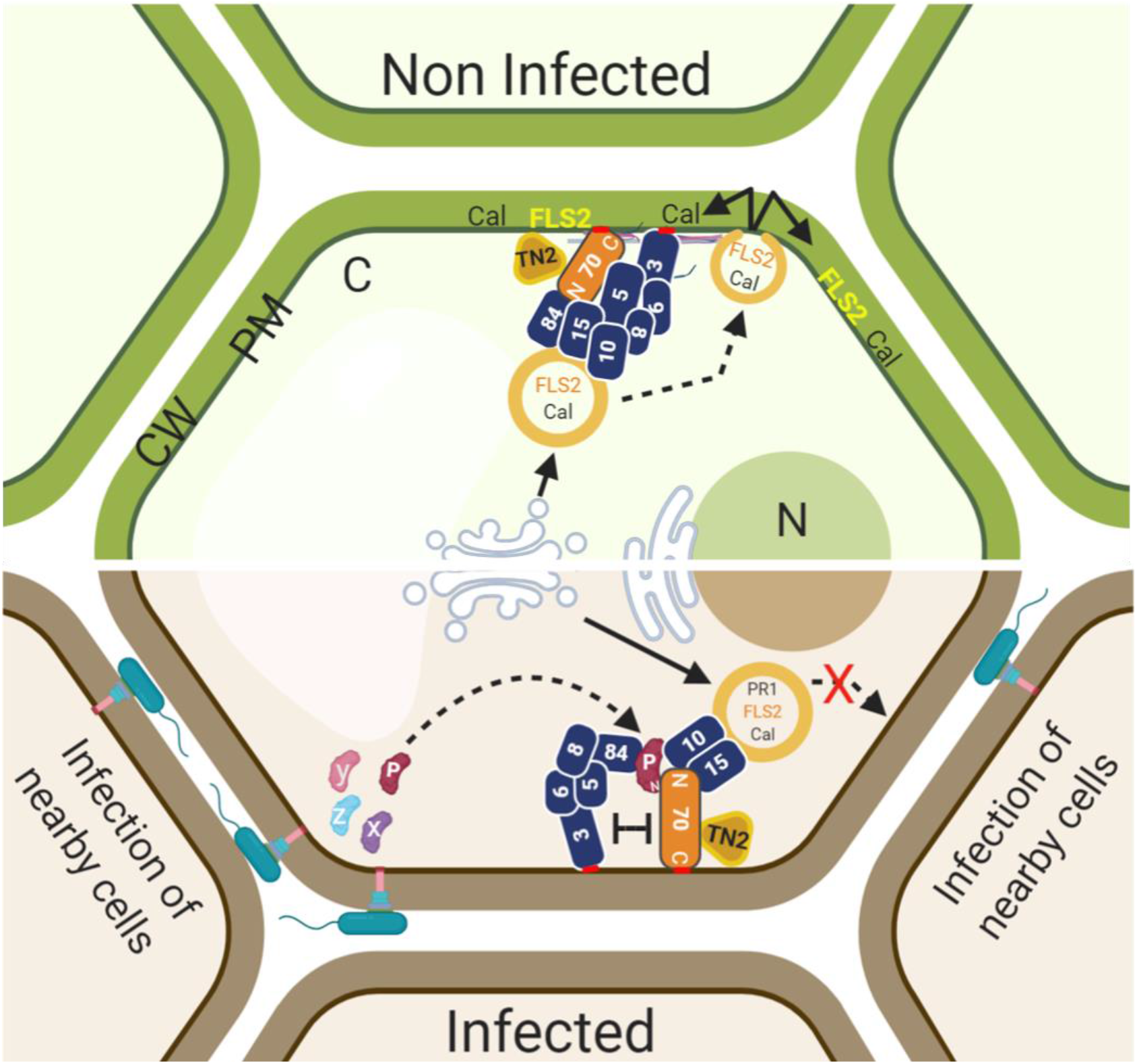
Schematical Abstract. *Xcc*XopP manipulates the exocytosis via associating with members of sub-complex II of exocyst, affecting its proper assembly with the sub-complex I.

## Conclusion

This work reveals a bacterial pathogenic strategy to hijack plant exocytosis though exocyst targeting to inhibit PTI, however, avoiding NLR-dependent immunity activation. Considering the contribution of XopP to *X. campestris* infection, it seems to play an important role in the development of bacterial black rot disease. As a consequence, the exocyst complex appears as a plant susceptibility component during *X. campestris* infection that could be a potential target for the development of strategies to generate resistance against bacterial diseases.

## Materials and Methods

### Plants

#### Nicotiana *species*

*Nicotiana benthamiana* and *Nicotiana tabacum* cv Petit Gerard and N34-4 plants, used in our study, were grown in greenhouse conditions.

#### Arabidopsis thaliana

*Arabidopsis thaliana* ecotype Columbia-0 (Col-0) was used in our study as wild type. Arabidopsis seeds were stratified at 4°C in water for 3 days before sowing in soil and then were grown in greenhouse conditions. To generate constructs for conditional expression of XopP effector from *Xanthomonas campestris* pv. *campestris* in Col-0 WT background, we inserted with Golden Gate cloning in series the native Rubisco promoter, the gene, a C-terminal mCherry epitope tag and NOS terminator into pICSL86955OD vector (Engler et al., 2008). The plants were transformed with the floral dip method (Clough and Bent, 1999). The transformants were selected with BASTA. In brief, plants were sprayed from a short distance with BASTA (Bayer, 120 ppm Glufosinate – ammonium) twice in a week and the survivals were screened through PCR.

#### Escherichia coli

*E. coli* (Stellar or DH10b strains) were grown on LB plates with the appropriate antibiotic at 37°C and kept at 4°C for up to two weeks. For liquid cultures, a bacterial scrape was inoculated in liquid LB with appropriate antibiotic and grown at 37°C with shaking at 200 rpm approximately.

#### Agrobacterium tumefasciens

*A.tumefasciens* (AGL1, C58C1 and GV3101 strains) were grown on LB plates with appropriate antibiotics at 28°C and kept at 4°C for up to two weeks. For liquid cultures, a bacterial scrape was inoculated in 5 ml LB with appropriate antibiotic and grown at 28°C with shaking at 200 rpm approximately.

### Bacterial strains

*Xcc*8004 WT, as well as 060E08 (mutant strain for XopP effector) were grown at 28°C in LB plates with appropriate antibiotics (Jiang et al., 2009).

### Cloning and Constructs

For all constructs, used in the study, we used Golden Gate (GG) cloning and Phusion-HF polymerase (NEB). All constructs generated were verified by Sanger sequencing. EXO70-ID was amplified from pDONR221_*Exo70*_C_C119, given from Prof. M. Moscou (TSL), with STOP codon using BsaI sites and cloned to pGBKT7-RFP GG compatible vector for Y2H assay. The EXO70-ID is a fusion of RGH2 NLR of *Hordeum vulgare* ssp. *vulgare* cv. Baronnesse. *AtExo70B1* and *AtExo70F1*, as there are intronless genes, were amplified from genomic DNA (from Col-0 plants) in three parts, domesticating the internal BsaI sites, either with STOP codon for Y2H assay or without STOP codon for tagging in plants and cloned to pBluescript GG compatible vector. For both analyses, GG reaction was performed with pGBKT7-RFP as a final vector for Y2H assay or pICH86988 with a C-terminal tag for plant assays. *AtExo70B2* was given to us in pUG41308 vector from Prof. Marco Trujillo (University of Freiburg) and subcloned to pGBKT7-RFP vector for Y2H assay. *ObExo70B1*_small was amplified from cDNA, in two parts, domesticating the internal BsaI site either with STOP codon for Y2H assay or without STOP codon for tagging in plants. The *ObExo70B1*_large was amplified from gDNA, first amplifying the whole ORF with the 5’ UTR and then performed nested PCR, with an internal FW primer. The PCR fragments were cloned firstly to pBluescript vector and then transferred in pGBKT7-RFP vector for Y2H assay or pICH86988 fused to a C-terminal epitope tag for plant assays. XopP was amplified from genomic DNA of *Xanthomonas campestris* pv. *campestris* ATCC33913 in two parts, domesticating the internal BsaI site with STOP codon for Y2H assay and without STOP codon for tagging in plants. *XopP1* and *XopP2* were amplified from genomic DNA of *Xanthomonas oryzae* pv. *oryzicola* WHRI 5234 (NCPPB 1585), its draft genome being recently sequenced (Michalopoulou et al., 2018). *Hlk3* was amplified from genomic DNA of *Ralstonia solanacearum* GMI1000, a kind donation of Prof. Nemo Peeters (INRA-CNRS). *AcXopP* was amplified from genomic DNA of *Acidovorax citrulli*, a kind donation of Prof. Goumas (HMU). All the above PCR fragments were cloned in pBluescript vector and then subcloned to pGADT7-RFP GG compatible vector for Y2H assay or to pICH86988 fused to a C-terminal tag for in planta assays.

Truncation forms of *At*EXO70B1 were generated with the help of Prof. Marco Trujillo, using *At*EXO70B2 as a template, due to high sequence similarity. Using pGBKT7-*At*EXO70B1 as a template, PCR fragments from 1 to 388 amino acids (AB), from 319 to 511 amino acids (BC) and from 389 to 625 amino acids (CD) were generated, including START and STOP codons and were cloned to pBluescript vector and then subcloned to pGBKT7-RFP for Y2H assays. Truncation forms of XopP were generated using Phyre2 bioinformatics program. Using pGADT7: XopP as a template, PCR fragments from 1 to 420 amino acids and from 421 to 733 amino acids, including START and STOP codons were amplified and cloned to pBluescript vector and then subcloned to pGADT7-RFP vector, for Y2H assay. For amplification of XopP minus the 100 first amino acids (XopP-100), an internal FW primer was used and was cloned from template pGADT7: XopP. AvrRps4 effector from *Pseudomonas syringae* pv. *pisi* was used from a previous study (Sarris et al., 2015).

Chp7 construct, used in the study, was given by Professor Jane Glazebrook (Lu et al., 2015). pMDC:spC7HPB was verified by restriction endonucleases digestion and *transferred* to AGL-1 *Agrobacterium tumefasciens* strain for plant assays (Lu et al., 2015). *XopQ* was amplified from genomic DNA of *Xanthomonas campestris* pv. *vesicatoria,* a kind donation of Prof. Goumas (HMU) and cloned to pICH86988 fused to a YFP C-terminal epitope tag. For the generation of PR1-RFP-GFP, PR1 secretion peptide was amplified from pMDC:spC7HPB construct using as a FW primer 5’ AAGGTCTCGccATGGGATTTGTTCTCTTTTCACAATTGC 3’ and as a RV primer 5’ AAGGTCTCACATTGAATTTTGGGCACGGCAAGAGTG. The PCR fragment was firstly cloned into pBluescript vector and sequence was verified by sequence analysis. The final construct was cloned into pICSL86900_OD under the expression of CamV 35S promoter and Nos terminator. FLS2-GFP construct, used in this study, was a kind donation of Dr. Katarzyna Rybak (LMU, Institute of Genetics), cloned in pCambia vector and transformed in GV3101 strain of *Agrobacterium tumefasciens*. *Tn2* NLR gene was amplified from *Arabidopsis thaliana Brassica napus* zs11 gDNAs and cloned to pBluescript vector and later subcloned to pICH86988, fused C-terminally to YFP epitope tag for in planta assays. The remaining exocyst components (*Sec3A*, *Sec5A*, *Sec6*, *Sec8*, *Sec10*, *Sec15A*, *Sec15B*, *Exo84B* and *Exo84C*) were amplified from *A. thaliana* cDNA and cloned into pBluescript vector and later to pGADT7 vector for Y2H assays or pICH86988 fused C-terminally to an epitope tag for in planta assays (stated in the text). DsRed-FYVE confocal marker was kindly given to us in a C58C1 *Agrobacterium* strain from Prof. František Baluška (University of Bonn). Plasma-Membrane-RFP, Tonoplast-RFP and Ubi10::mCherry::Atg8A::Alli mCherry confocal markers were kindly given to us in AGL-1 (but for Atg8a which was in GV3101) *Agrobacterium* strains from Prof. Yasin Dagdas (Gregor Mendel Institute of molecular plant biology).

### Phylogenetic analysis of XopP

In KEGG, using *Xanthomonas campestris* pv. *campestris* XopP (XCC_1247) in search term, we identified the orthologs from various pathogens, including *Xanthomonas* species, *Ralstonia* and *Acidovorax.* The annotated protein sequences of 73 orthologs were manually downloaded from KEGG and two of them were recovered from sequence analysis of draft genome annotation of *Xanthomonas oryzae* pv. *oryzicola* WHRI 5234 (NCPPB 1585) (Michalopoulou et al., 2018). Identity percentage of orthologs used did not drop down from 24,4%. Moreover, in order to confirm that the resultant orthologs are indeed homologous to Xcc_1247, we did reverse Blastp in NCBI for each of the 73 protein sequences. From this search, we excluded nine protein sequences. All 67 proteins were aligned via MEGA5. MEGA5 was also used to construct the maximum likelihood phylogenetic tree using the Poisson correction model (Tamura et al., 2011; ZUCKERKANDL and PAULING, 1965).

### Y2H and Y3H assays

The vectors used for Y2H assay were pGBKT7-RFP and pGADT7-RFP, which were GG compatible, kindly given to us from Dr. Marc Youles (TSL). RFP is a dual selection marker, along with the antibiotic resistance gene, being utilized for inserting the gene of interest. All plant proteins were cloned as baits (in pGBKT7 vector) and bacterial effectors were cloned as preys (in pGADT7 vector), otherwise being stated. A standard two-hybrid screen was carried out with the yeast reporter strain PJ694a (or otherwise stated where we used AH109), using all the necessary negative controls (bait with empty pGADT7-RFP, prey with empty pGBKT7-RFP and empty pGBKT7-RFP and pGADT7-RFP) for each transformation pair. As a positive control, a known interaction was used from a previous study (*Sarris et al.,* 2015). The selection of the plasmids transformation was and yeast-two hybrid analysis was performed with selection of the auxotrophic markers leucine, tryptophan and adenine, respectively. Accordingly, the vectors used for Y3H assay were pBridge, kindly given to us by Prof. Subba Rao Gangi Setty (Indian Institute of Science Bangalore) and pGADT7-RFP vectors. EXO70B1 and XopP were cloned in MCS I and II sites of pBridge, respectively. AH109 yeast strain was transformed with both vectors and the two-hybrid analysis was performed with selection of the auxotrophic marker adenine with supplementation of 1mM Methionine in the medium, for suppressing XopP expression. The three hybrid analysis was performed in medium lacking leucine, tryptophan, adenine and methionine. All yeast strains left to grow in 30°C.

### Subcellular co-/localizations and BiFC

*N. benthamiana* was infiltrated with *Agrobacterium* strains at OD_600_ = 0.5 expressing indicated constructs. Confocal markers for co-localization assays were expressed at OD600 = 0.4. For subcellular co-/localizations and BiFC, leaf discs were analyzed 3 dpi. All images were captured in a Leica SP8 confocal microscope (facility of IMBB). For subcellular localizations, all indicated constructs were fused to a C-terminal eCFP, YFP or mCherry epitope tags, unless for the confocal markers which were given to us already cloned with DsRed, mCherry or RFP epitope tags. For BiFC, all indicated constructs were fused to a C-terminal nVENUS or nCERULEAN or cCFP epitope tags. All samples were imaged with a 40x water objective, otherwise marked. The confocal images were processed with ImageJ. Excitation of GFP was made using the 488 nm of Argon laser and emission of spectra was collected at 496-520 nm. Excitation of YFP was made using the 514 nm of Argon laser and emission of spectra was collected at 520-550 nm. Excitation of mCherry, *Ds*Red and RFP was made using the 561 nm laser and emission of spectra was collected at 593-628, 573-616 or 573-620 nm, respectively. Excitation of eCFP was made at 405 nm and emission of spectra was collected at 463-487 nm. Chlorophyll was detected between 653 and 676 nm and excited from 561 nm. Gain was at 100%, otherwise marked. Quantification of GFP signal from infiltrations with FLS2-GFP was processed with Image J software, as described in https://theolb.readthedocs.io/en/latest/imaging/measuring-cell-fluorescence-using-imagej.html.

### HR assay

For HR assay with PR1sp-Chp7 in presence or absence of XopP *N. tabacum* cv. Petit Gerard were infiltrated with *Agrobacterium* C58C1 or AGL-1 strains at OD_600_ = 0.1 for PR1sp-Chp7 and OD_600_ = 0.4 for GUS and XopP. For HR assay with *At*- or *Bn*TN2 constructs, *N. tabacum* cv. N34-4 and Petit Gerard were infiltrated with *Agrobacterium* C58C1 strain of TN2, EXO70B1 or XopP at OD_600_ = 0.5 (equimolar ratio).

### Callose deposition

Leaves from 4-week-old *Arabidopsis* plants were infiltrated with 20 μM of flg22 in 10 mM of MgCl_2_ or mock solution and removed 12 h after infiltration. Callose staining, image acquisition, and processing were carried out as described in (Lin Jin, 2017) with a change in destaining. In brief, clearing leaves from cholorophyll was carried out with acetic acid and ethanol in a 1:3 ratio overnight (12 hours). Callose visualization was made in confocal SP8 microscope in a 10x dry objective with excitation from 405 nm and emission between 490 and 530 nm. Images were processed as described in (Lin Jin, 2017) with Image J software and a graph was created depicting Callose deposits/mm2, from 6 pictures per genotype and treatment. Some samples with extreme values were removed from the quantification.

### Protein extraction and Co-immunoprecipitation

*N. benthamiana* was infiltrated with Agrobacterium strains at OD_600_ = 0.5 expressing indicated constructs. Agro-infiltrated leaves were grinded 72 hpi with liquid nitrogen and proteins were extracted in GTEN buffer (10% glycerol, 25 mM Tris-Cl pH7.5, 1 mM EDTA, 150 mM NaCl) supplemented freshly with 10mM DTT, protease inhibitor cocktail (Calbiochem) and 0,15% v/v NP-40. After 10 min centrifugation at 4°C, supernatant was collected in clean microcentrifuge tube and incubated 2 h at 4°C with a-flag mouse antibody (Sigma), following 1 h incubation at 4°C with PureProteomeProtein A/G Magnetic Beads Mix (Millipore). Beads were washed with GTEN buffer, before adding 1x SDS loading dye and boiling the samples for 5 minutes. Total protein from yeast transformations was extracted according to (Horvath et al., 1994).

### SDS PAGE and Immunoblotting

Protein samples from plant leaves were separated at 8% SDS PAGE, electro-blotted at PVDF membrane (Millipore), then blocked for 1 h with 5% w/v milk in TBS and incubated for 1 h with primary antibodies a-flag 1:4000 (Sigma) or a-myc 1:4000 (Cell Signalling). Membranes were then incubated for 1 h with secondary antibody a-mouse HRP conjugate 1:10000 (Cell Signalling) and chemiluminescent substrate ultra (Millipore) was applied for 5 min before camera detection in Sapphire Biomolecular Imager (Azure Biosystems). Total protein lysate from yeast transformations were separated in 10% or 12% SDS PAGE, electro-blotted at PVDF membrane (Millipore), then blocked for 90 minutes with 5% w/v milk in TBS supplemented with 0,05% Tween-20 and incubated for 1 h with the primary antibody a-Gal4-DBD 1:500 (sc-510, Santa Cruz Biotechnology). The rest procedure is similar as in proteins extracted from plant leaves.

### Protein expression conditions in *E.coli*

Different fragments of XopP protein were produced (Fig. S2A), several of which were insoluble. We determined, via multiple alignment of XopP homologues in *Xanthomonas* genus, that the only soluble construct, until now, is the fragment lacking the first 100 N-terminal residues (molecular weight 71.092 Da). In case of EXO70B1, its full length of 624 amino acid residues (molecular weight 70.639 Da) was a soluble construct.

*XopP-100* gene fragment was cloned and inserted into the pET26b (ColE1 plasmids) vector carrying a C-terminal 6-His epitope tag and transformed into the *E.coli* strain BL21. A sufficient amount of soluble protein was obtained after induction using the following conditions: Cells were grown in LB medium containing 30 μg/ml kanamycin until an OD_600_ value of 0.6 was reached. The culture was induced with 0.4 mM IPTG and left to grow at 20°C overnight. The cell paste was re-suspended in 100 ml lysis buffer containing 25 mM Tris pH=8.0, 300 mM NaCl, 5 mM imidazole and 15 mM β-mercaptoethanol, and homogenized. After adding protease inhibitors (20 μg/ml leupeptin, 1 mM PMSF and 150 μg/ml benzamidine), the solution was sonicated for 5 min. The precipitate was subsequently removed by centrifugation at 12,000 rpm at 4°C for 45 min. Purification was carried out using His-tag affinity chromatography at 4°C with a 8-ml Ni-NTA Qiagen column pre-equilibrated in lysis buffer and initially washed stepwise with 10, 20 and 30 mM imidazole. With a subsequent increase in imidazole concentration, the protein started eluting at 100 mM imidazole. Fractions containing the protein were dialyzed against the storage buffer containing 25 mM Tris pH=8, 100 mM NaCl and 10 mM β-mercaptoethanol and concentrated to approximately 2.8 mg/ml, for subsequent crystallization experiments. In case of *Exo70B1*, the full length gene was cloned and inserted into the LIC 1.10 vector carrying a N-terminal 6-His-SUMO3 tag, and transformed into the *E.coli* strain Rosetta™ (DE3). The same protocol was followed and the fractions containing the protein were dialyzed against the storage buffer (25 mM Tris pH=8, 100 mM NaCl, 10 mM β-mercaptoethanol and 0.5 nM SENP-2 protease). Subsequently, a reverse purification protocol was carried out using His-tag affinity chromatography, and the fractions containing the protein were concentrated to approximately 8.5 mg/ml.

### Field-Emission Scanning Electron Microscopy (FE-SEM)

The XopP-100 fragment was purified by His-tag affinity chromatography. Fractions containing the protein were dialyzed against the storage buffer in absence of β-mercaptoethanol overnight. The gel-like form that came up, was deposited on a covered glass and air-dried overnight. The sample was then covered with 10 nm of Au/Pd sputtering and observed using a JEOL JEM-2100 transmission electron microscope at 20 kV.

### Transmission Electron Microscopy (TEM)

5 μl of XopP-100 sample was deposited onto a formvar/carbon-coated electron microscopy grid for 2 min. After removing the excess with filter paper, 2% w/v uranyl acetate was used in order the sample to be stained, for 2 min. The observations were conducted with a JEOL JEM-2100 transmission electron microscope at 80 kV

## Acknowledgments

V.A.M was supported by the Hellenic Foundation for Research and Innovation (HFRI) and the General Secretariat for Research and Technology (GSRT), under the HFRI PhD Fellowship grant (GA. no. 4776) and by the internal PhD supporting scheme of IMBB-FORTH. We thank especially Prof. Marco Trujillo for his generous donation of *At*EXO70B2 construct as well as for his advice on truncations construction; Prof. M. Moscou for his kind donation of some of the Integrated Domains used in the reseach; Dr. Katarzyna Rybak for the kind donation of FLS2-GFP construct; Prof. František Baluška for his kind donation of DsRed-FYVE confocal marker; Dr. Yasin Dagdas for his generous donation of all the other confocal markers used in the research; Mr Marc Youles and the Synthetic Biology Group of TSL for the kind donation of Golden-Gate compatible yeast-2-hybrid vectors; Prof. Nemo Peeters for his kind donation of *Rs* GMI1000; Prof. Dimitris Goumas for his generous donation of *Xcv*; Prof. Subba Rao Gangi Setty for his kind donation of pBridge vector and lastly undergraduate student Aspa Papanikolaou for her kind contribution in some of the Y2H assays.

## Authors Contributions

V.A.M. carried out most of the experiments; K.K., D.K., M.K. and P.H.N.C. purified recombinant proteins and performed *in vitro* interaction assays; G.M. performed additional experiments and contributed to organizing some of the experiments in the Lab. V.A.M. performed *Arabidopsis* transformation; P.F.S. designed the research and supervised the project; P.F.S. and V.A.M. initiated the project; V.A.M. P.F.S. and J.D.G.J. analysed the data/results. P.F.S., V.A.M. and J.D.G.J. wrote the manuscript. All authors have read and approved the manuscript.

