## Supplementary Figures for "The host exocyst complex is targeted by a conserved bacterial type III effector protein that promotes virulence"

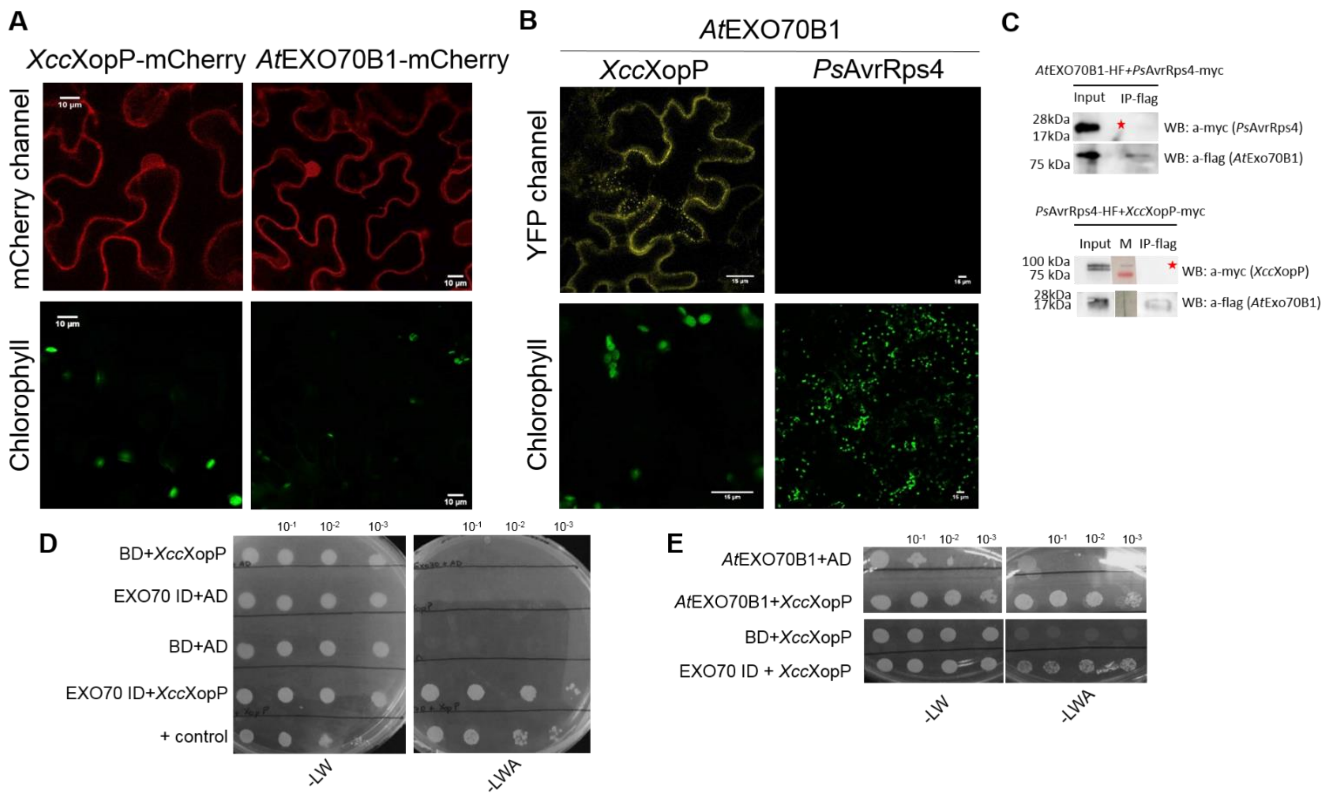


**Figure S1. *Xcc*XopP and *At*EXO70B1 co-localization assays. EXO70-ID and *At*Exo70B1 interact specifically with *Xcc*XopP effector. EXO70B1 and *Xcc*XopP do not interact with AvrRps4 (negative control), in a co-IP assay.**

**A.** *Xcc*XopP and *At*EXO70B1 localization. The effector and the plant proteins are fused C-terminally with mCherry epitope tag and the pictures are taken in confocal microscopy 3dpi in *N. benthamiana* leaves.

**B.** *At*EXO70B1 interacts with *Xcc*XopP effector (left lanes) but not interact with AvrRps4 effector (right lanes) in a BiFC assay. *At*Exo70B1 is C-terminally fused with nVENUS tag and both effectors with cCFP tag. YFP signal is observed in confocal microscopy 3dpi in *N. benthamiana* leaves.

**C.** *At*EXO70B1 and *Xcc*XopP do not interact with AvrRps4 effector (marked with red asterisks), used as negative control, in a co-IP assay.

**D, E**. Y2H assay of EXO70-ID (D) and *At*EXO70B1 (E) in pGBKT7 and *Xcc*XopP in pGADT7, as well as, their negative controls. There have been made four serial dilutions. Pictures have been taken in the third day of yeast growth in plates lacking leucine, tryptophan and/or adenine.

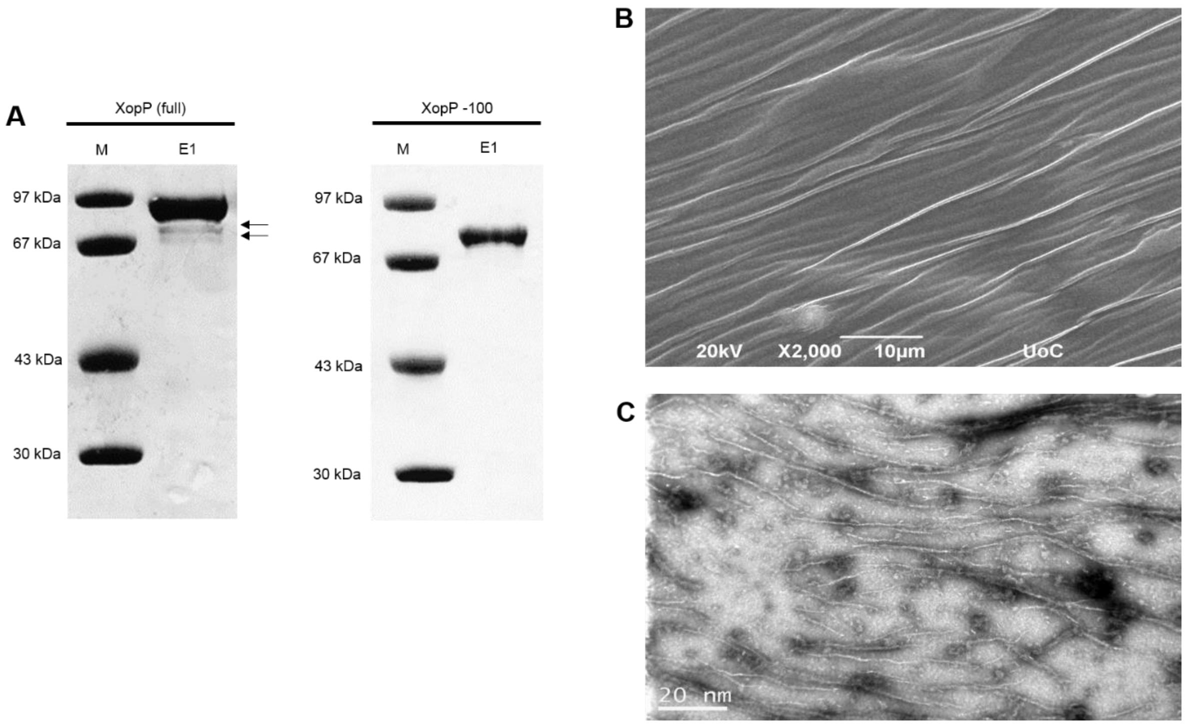


**Figure S2. *Xcc*XopP expression pattern and formation of fibrils.**

**A.** Elution fractions of *Xcc*XopP in a Sodium Dodecyl Sulfate–Polyacrylamide Gel electrophoresis (SDS-PAGE) of a 10% acrylamide gel (A) and *Xcc*XopP-100 (B) after His-tag affinity chromatography with Ni-NTA Qiagen column, reveals severe cleavage only in the case of full length *Xcc*XopP protein (black arrows).

**B.** Visualization of field-emission SEM (FE-SEM) picture of the self-assembled *Xcc*XopP-100 gel-like formation after overnight dialysis in storage buffer in absence of β-mercaptoethanol. Morphology of fibrils is observed. FE-SEM scale bar = 10 μm. D. Visualization of TEM micrographs of the *Xcc*XopP-100 fibrils stained negatively with 2% uranyl acetate, after overnight dialysis in storage buffer in absence of β-mercaptoethanol. Scale bar = 20 nm.

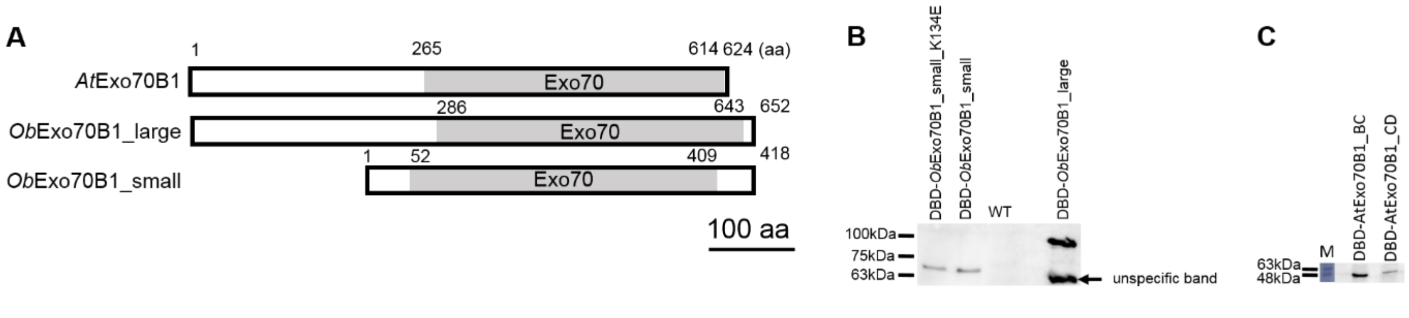


**Figure S3. Large and small versions of *Ob*Exo70B1 compared with *At*Exo70B1. Expression of small *Ob*EXO70B1 and its mutant K134E.**

**A.** Schematic representations of *At*Exo70B1 and its homolog *Ob*Exo70B1 (large and small version). Only large version was annotated in NCBI. Small version has 234 missing amino acids from the N-terminal site and it was detected by our group. With grey appears the Exo70 catalytic domain in each homolog, as it was pointed out by SMART. Bar = 100 aa.

**B.** Western blot analysis of WT *Ob*EXO70B1_small and _large versions and the mutant K134E_small in an SDS PAGE (8% acrylamide gel). The proteins are fused N-terminally with DBD of Gal4 transcription factor and are of 66 kDa (small construct) and 90 kDa (large construct).

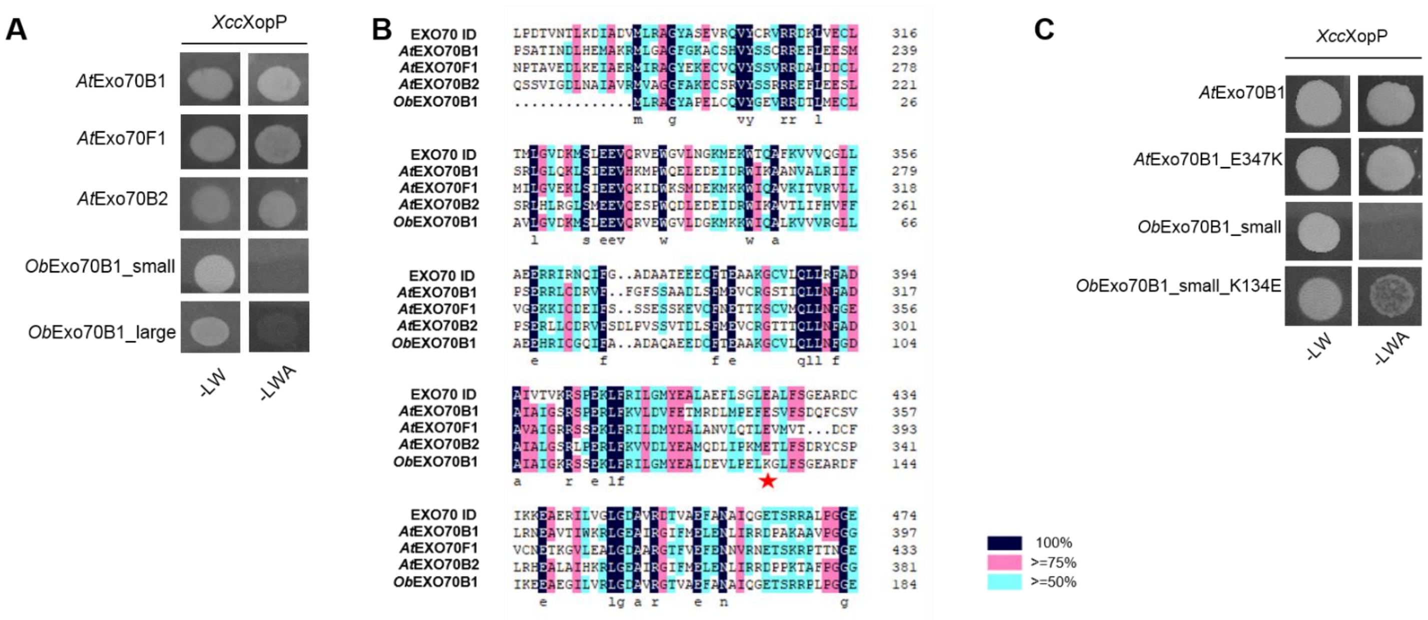


**Figure S4. *Xcc*XopP associates with *Arabidopsis* EXO70B2 and EXO70F1 but not with the EXO70B1 from monocots. Also, E347 is essential for *At*EXO70B1 interaction with *Xcc*XopP.**

**A.** Y2H screening between *Xcc*XopP and *At*Exo70B1, *At*Exo70F1, *At*Exo70B2 and *Ob*Exo70B1_small and _large versions. All of the plant proteins, but *Ob*Exo70B1 (both versions) interacted with the effector.

**B.** Representation of protein alignment between all interacting and no-interacting proteins, including the EXO70-ID is presented. The area of alignment is focused to the AB domain of *At*Exo70B1 that interacts with *Xcc*XopP. K134 of *Ob*EXO70B1_small, marked with a red star, is the only pointed amino acid that appears to be conserved between the interacting homologs and mutated in the non interacting *Ob*Exo70B1.

**C.** Y2H screening between *Xcc*XopP with *At*EXO70B1, *At*EXO70B1$^{E347K}$, *Ob*EXO70B1_small or *Ob*EXO70B1_small$^{K134E}$. K134E seems to be capable of gaining the interaction between *Ob*EXO70B1_small$^{K134E}$ and *Xcc*XopP, however, it is not essential to abolishing the *At*EXO70B1$^{E347K}$/*Xcc*XopP interaction.

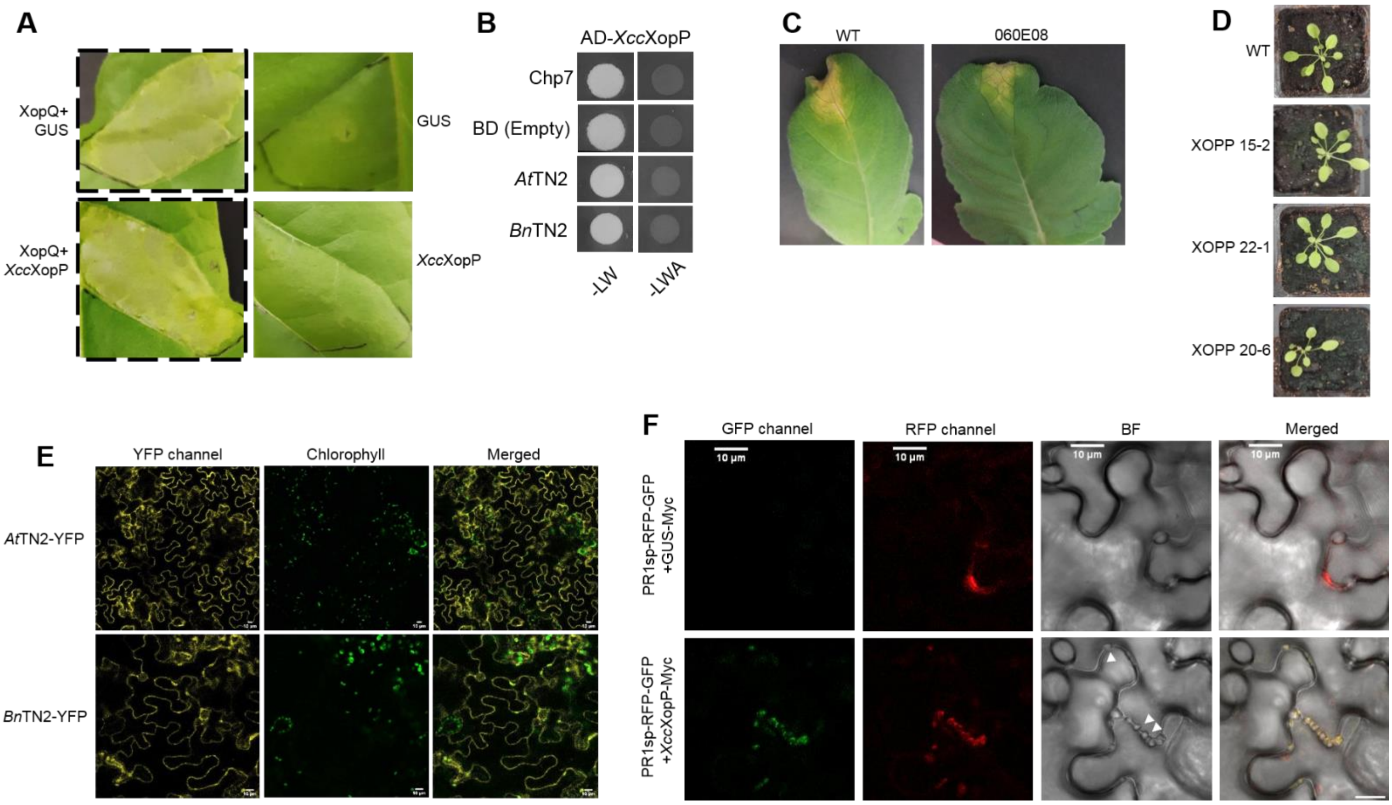


**Figure S5. *Xcc*XopP is not a general cytoplasmic HR inhibitor nor interacts with Chp7 effector and TN2 from *Arabidopsis thaliana* and *Brassica napus*. Pathogenicity of the mutant XopP *Xcc*8004 strain (060E08) is reduced in *Raphanus sativus*. Expression of *Arabidopsis thaliana* and *Brassica napus* TN2. Inhibition of PR1A exocytosis.**

**A.** The co-expression of *Xcc*XopP with XopQ effector, from *Xanthomonas campestris* pv. *vesicatoria* (left lower lane), did not inhibit the cytoplasmic HR induction caused by the XopQ (left upper lane). Expression of GUS or *Xcc*XopP itself does not induce any HR (right lanes).

**B.** *Xcc*XopP does not interact with Chp7 effector and *At*-TN2 and *Bn*-TN2 NLRs in a Y2H screen, as it is observed in SC-LWA medium (right lanes).

**C.** Pathogenicity test of WT and *Xcc*XopP mutant strain (060E08) *Xcc*8004 w.t. in *Raphanus sativus.* A reduced virulence is shown in the case of *Xcc*XopP mutant as it was previously reported. The picture is taken in the 10$^{th}$ days post infection and the experiment was repeated three times with similar results.

**D.** Expression of *At*-TN2 and *Bn*-TN2 fused C-terminally to YFP epitope tag, in *Nicotiana benthamiana* leaves. The pictures are taken in a confocal microscope 3 dpi. Bars = 10μm.

**E.** Visualization of PR1sp-RFP-GFP inhibition of exocytosis by *Xcc*XopP in confocal microscope. The construct was co-expressed with either *Xcc*XopP-myc or GUS fused C-terminally to myc (as a negative control) in *N. benthamiana* leaves and images were taken 3dpi. Bars = 10 μm.

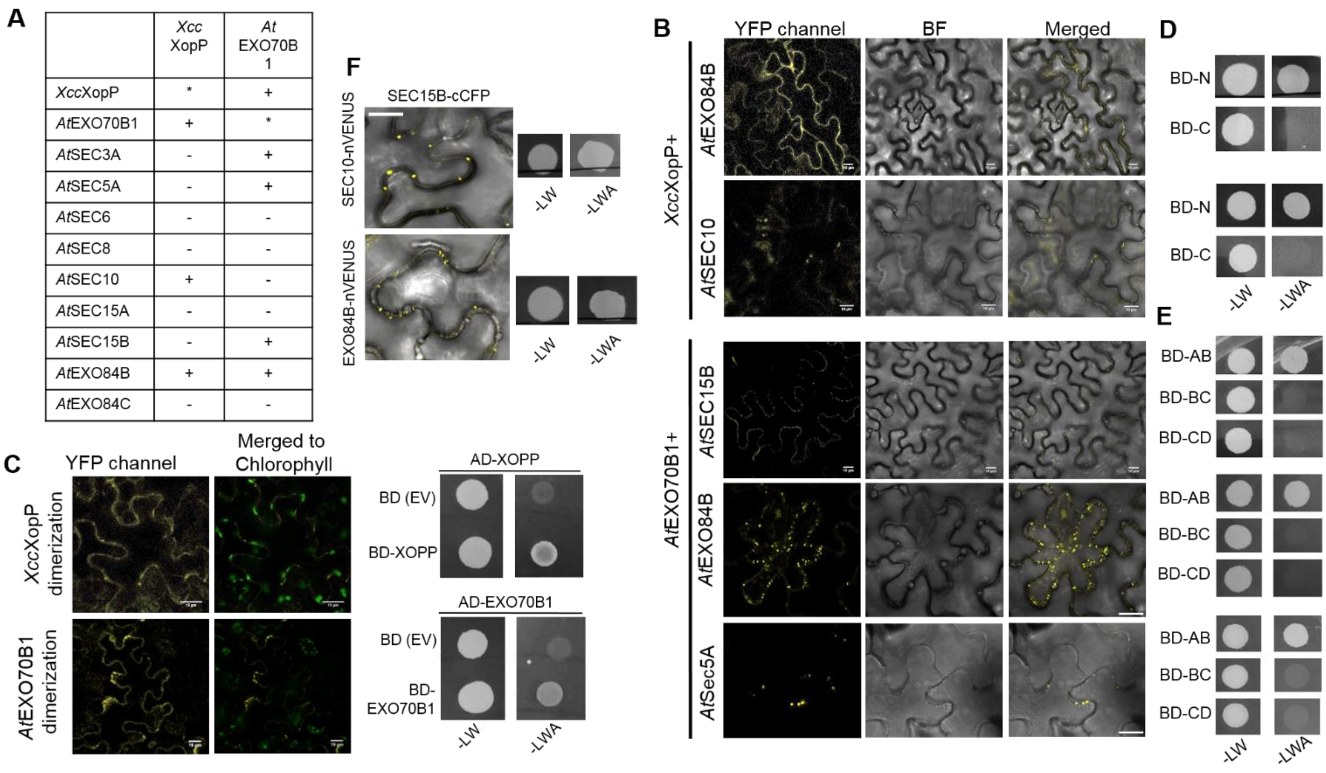


**Figure S6. *Xcc*XopP interacts with more than one exocyst components and with itself. Also, EXO70B1 interacts with SEC15B, EXO84B, SEC5A and with itself.**

**A.** Binary interactions of *Xcc*XopP and EXO70B1 with eight components of the exocyst complex, including self interactions, in a Y2H screening. The components listed in the vertical axis were cloned as baits and those in horizontal axis were cloned as preys and transformed in PJ69a strain of *Saccharomyces cerevisiae.* +: positive interactions (with lower size weak interactions are presented, -: negative interactions, *: positive dimerization.

**B.** Validation of the interactions *in planta*, using BiFC assays. EXO84B, SEC10 and EXO70B1 were fused C-terminally with nVENUS and *Xcc*XopP, SEC15B, EXO84B and SEC5A with cCFP epitope tags and transiently co-expressed in *N.¨benthamiana* leaves. The pictures are taken 3dpi. Bars = 10 μm.

**C.** Dimerization of *Xcc*XopP and *At*EXO70B1 using BiFC and Y2H assays. *Xcc*XopP and EXO70B1 were cloned C-terminally with both nVENUS and cCFP epitope tags and transiently co-expressed in *N. benthamiana* leaves. 3dpi images were taken in a confocal microscope. Bars = 15 μm. For the Y2H screening, EXO70B1 and *Xcc*XopP were cloned simultaneously as baits and preys and each protein was tested for dimerization in *S. cerevisiae.*

**D.** Interactions of SEC10 and EXO84B with *Xcc*XopP N- and C-terminal regions in Y2H assays. Both proteins interact with the N-terminus of the effector, as visualized in medium lacking LWA.

**E.** Interactions of SEC15B, EXO84B and SEC5A with *At*EXO70B1 truncations in Y2H assays. All proteins interact with the N-terminus of *At*EXO70B1 (AB truncation), as visualized in medium lacking LWA.

**F.** SEC10 and EXO84B interact with SEC15B in BiFC and Y2H analyses. For the BiFC assay, *sec10* and *exo84b* were fused to nVENUS C-terminal tag and *sec15b* to cCFP C-terminal tag and co-infiltrated in *N. benthamiana* leaves. Three dpi, YFP signals were observed in confocal microscopy. Bars 10 μm. For Y2H analysis, *sec10* and *exo84b* were cloned as baits and *sec15b* as prey. Each pair of constructs was co-transformed to PJ69 yeast strain and 4 days later, yeast growth was observed in medium lacking leucine, tryptophan and adenine.

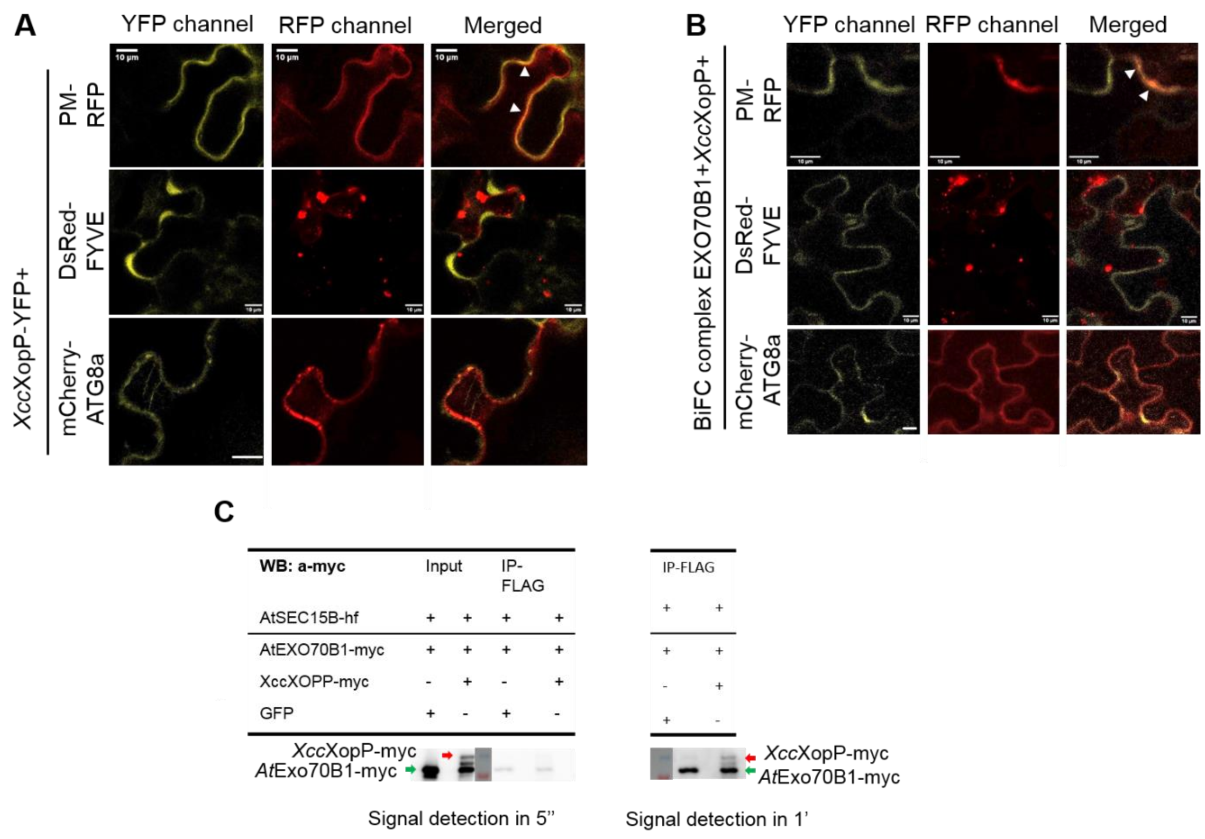


**Figure S7. *Xcc*XopP and in the EXO70B1/*Xcc*XopP complex localize to the plasma membrane.**

**A, B.** co-localization assays of *Xcc*XopP (A) and EXO70B1/*Xcc*XopP complex (B) with compartmental markers. Plasma Membrane (PM) is fused c-terminally to RFP, phosphatidyl-inositol-3-phosphate(PI(3)P)-enriched early/late endosomal compartmental marker FYVE is N-terminally fused to *Ds*Red, autophagosome marker ATG8a is fused N-terminally with mCherry epitope tags. The effector itself, as well as, the EXO70B1/*Xcc*XopP complex co-localize with the PM marker (white arrowheads). All the above markers were co-expressed with either *Xcc*XopP fused C-terminally to YFP, or the EXO70B1/*Xcc*XopP complex, fused c-terminally to nVENUS and cCFP epitope tags, respectively, IN *N. benthamiana* leaves and confocal images were visualized 3dpi. Bars = 10 µm.

**C.** *Xcc*XopP did not inhibit EXO70B1-SEC15B interaction in a triple co-IP assay. *XccXopP* and *AtExo70B1* were fused to -myc C-terminal tag and *AtSec15B* to -HF C-terminal tag and triple agro-infiltration of equimolar ratio was performed in *N. benthamiana* leaves. Three dpi the tissue was grinded and total proteins were extracted. Triple co-IP was performed with FLAG-beads and sequentially western blot analysis was performed with anti-myc for signal detection of EXO70B1-myc in absence and presence of *Xcc*XopP. The protein levels were not affected in presence of *Xcc*XopP, as well as the effector was also co-immunoprecipitated via *At*EXO70B1, as it appears in prolonged exposure time.
